# An Engineered Living Material with pro-angiogenic activity inducible by near-infrared light

**DOI:** 10.1101/2025.11.14.688407

**Authors:** Anwesha Chatterjee, Stefanie S.M. Meier, Sara Trujillo, Andreas Möglich, Shrikrishnan Sankaran

## Abstract

Impaired angiogenesis is a central barrier in the treatment of chronic and deep tissue wounds, preventing progression through the normal healing cascade. While the combination of near-infrared (NIR) photobiomodulation and pro-angiogenic growth factors has shown synergistic therapeutic benefit, the clinical translation of growth factor therapy is hindered by high cost, instability and the need for localized dosing to avoid aberrant vasculature. Peptidomimetics such as the VEGF-derived QK peptide offer a more stable and predictable alternative, but still require a means for localized, tunable presentation. Here, we establish an engineered living material based delivery system that responds to clinically relevant NIR light to produce and releases a QK-Fusion protein directly at the target site. The probiotic *Escherichia coli* Nissle 1917 was engineered with an 800 nm-responsive optogenetic circuit and encapsulated within an optimized alginate core–shell hydrogel that ensures biocontainment while allowing controlled outward diffusion of the secreted peptide. The released peptide remains non-cytotoxic and capable of binding extracellular matrix analogs and promoting the formation of organized, branched capillary-like networks in endothelial cultures. We thus establish a strategy for developing engineered living materials towards remote-controlled angiogenic stimulation.

## 1. Introduction

Healing of complex dermal defects remains a major clinical challenge because the native cascade of hemostasis, inflammation, proliferation and remodeling is often derailed by infection, metabolic disorders or poor vascularization. Photobiomodulation with red and near-infrared (NIR, 650–950 nm) light has emerged as a non-invasive strategy to re-establish this cascade: NIR light penetrates several millimeters through tissue, boosts mitochondrial ATP production, dampens excessive inflammation and accelerates collagen deposition, all while avoiding thermal damage [^1^]. In pre-clinical rodent and porcine models, pulsed 810 nm or 830 nm lasers shortened closure times and markedly increased granulation tissue and capillary density [^2^,^3^], and first-in-human studies now report faster epithelialization of venous leg ulcers [^4^].

Angiogenesis is a decisive bottleneck in tissue regeneration, and several groups therefore pair NIR illumination with exogenous pro-angiogenic cues. For example, topical recombinant Epidermal Growth Factor (EGF) delivered alongside 830-nm light closed a deep burn in a human patient within five treatments [^5^] and, in a delayed tooth-replantation model, adjunctive basic Fibroblast Growth Factor (FGF) plus 808-nm light administration improved periodontal regeneration and curbed root resorption compared with either therapy alone [^6^]. Despite this promise, translation of recombinant growth factors is hampered by two long-standing obstacles. First, their picomolar potency means that even modest overdosing induces aberrant, leaky vasculature, oedema or tumor-like lesions, necessitating tight local concentration control [^7^]. Second, large-scale production, purification and cold-chain logistics render full-length proteins prohibitively expensive [^8^]. Furthermore, these growth factors are susceptible to enzymatic degradation and hydrolysis in the body, thereby requiring large initial doses [^9^]. Vascular Endothelial Growth Factor (VEGF), for instance, costs orders of magnitude more per milligram than standard biologics yet shows a narrow therapeutic window and off-target effects when chronically over-expressed [^10^].

Material scientists and protein engineers have tackled these hurdles along two complementary lines. First, smart delivery vehicles have been advanced to control growth factor dosing [^11,12^]. These vehicles comprise of micro-/nanoparticles, core–shell hydrogels, photo-responsive matrices, etc. and provide on-demand, localized release while minimizing systemic spill-over. In one study, for instance, gold nanorod-conjugated liposomes embedded in collagen hydrogels enabled the release of both VEGF and platelet-derived growth factor (PDGF) at different NIR wavelengths [^13^]. In a different strategy, injectable black phosphorous nanosheet hydrogels released molecules upon exposure to 808 nm NIR light [^14^]. Furthermore, hydrogels containing MXenes (2D metal carbides with photothermal properties) functioned as effective photothermal platforms, releasing growth factors like VEGF or EGF under controlled conditions by transforming NIR light into localized heat [^15,16^]. Second, peptidomimetics - small fragments of the growth factors - are being developed to reduce the cost of growth factor therapies, as they are smaller and cheaper to produce. These synthetic analogs simulate either the key functional or structural domains of the native proteins while providing high stability, reduced immunogenicity and improved bioavailability [^17^]. Some examples include the SPPEPS sequence (TGF-β mimic) for promoting chondrogenesis [^18^], single-chain tandem macrocyclic peptides (STaMPtides) as Hepatocyte Growth Factor (HGF) receptor agonists [^19^], and VEGF peptide mimetics, which supports vascularization [^20,21^]. Among VEGF peptidomimetics, the QK peptide—derived from the α-helical region of VEGF_165_—has been widely studied for its ability to stimulate endothelial cell proliferation and capillary formation. Similar to native VEGF, QK demonstrates better angiogenic activity when immobilized onto the extracellular matrix, increasing its chemotactic activity [^20^].

Engineered living materials (ELMs) now promise to unite these advances in a single platform. By embedding genetically programmed microbes inside a biocompatible matrix, ELMs enable continuous, self-renewing synthesis of therapeutic payloads whose timing and dose are set by user-defined stimuli. We recently demonstrated the concept by entrapping blue-light-responsive *E. coli* in Pluronic F127 hydrogels to secrete collagen-binding QK on cue. Subsequent reports using alginate or gelatin matrices, probiotic lactobacilli and diverse inducible triggers underline the versatility and translational potential of ELMs [^22,23^].

Here we build on these foundations and report an NIR-light-regulated, pro-angiogenic ELM tailored for deep-tissue wounds. Our system harnesses the clinically proven probiotic *Escherichia coli* Nissle 1917, endowed with our recently developed NIR-light-responsive optogenetic circuit, to synthesize and secrete a QK-based fusion protein only when illuminated with 800 nm light. The bacteria are confined within core–shell alginate beads whose nanoporous shell prevents cell escape yet allows protein diffusion. We show that protein release from these ELMs is remotely switchable and tunable using NIR light, and that the secreted QK protein is functional, non-cytotoxic, and sufficient to induce angiogenic network formation in 2D cultures of vascular endothelial cells.

## 2. Results and Discussion

### 2.1. Engineering *E.coli* Nissle 1917 for light-responsive secretion

For this study, we used our recently developed optogenetic circuit *Av*NIRusk that controls bacterial gene expression by NIR light [^24^]. The light signal is translated by an engineered two-component system into a gene expression output. Specifically, the first component is a chimeric histidine kinase consisting of the light sensor module from the bacterial phytochrome *Agrobacterium vitis* (*Av*PCM) and the catalytic domains of the histidine kinase FixL from *Bradyrhizobium japonicum*. In the presence of NIR light, the chimeric histidine kinase phosphorylates the second component, the response regulator FixJ from *B. japonicum*. In its phosphorylated state, FixJ binds to the FixK2 promoter and initiates gene expression. All these components and the gene of interest are encoded on the single *Av*NIRusk plasmid (**Figure 1a**). As an initial validation, we tested the expression of the fluorescent reporter *Ds*Red in *E. coli* Nissle 1917 (EcN), which was upregulated a hundred-fold under NIR light with minimal basal expression (Figure S1a-c). We further performed a dose-response experiment using *Ds*Red fluorescence as output with varying NIR intensities to determine the optimal power level for the maximal activation of the system (Figure S1d).

**Figure 1.**
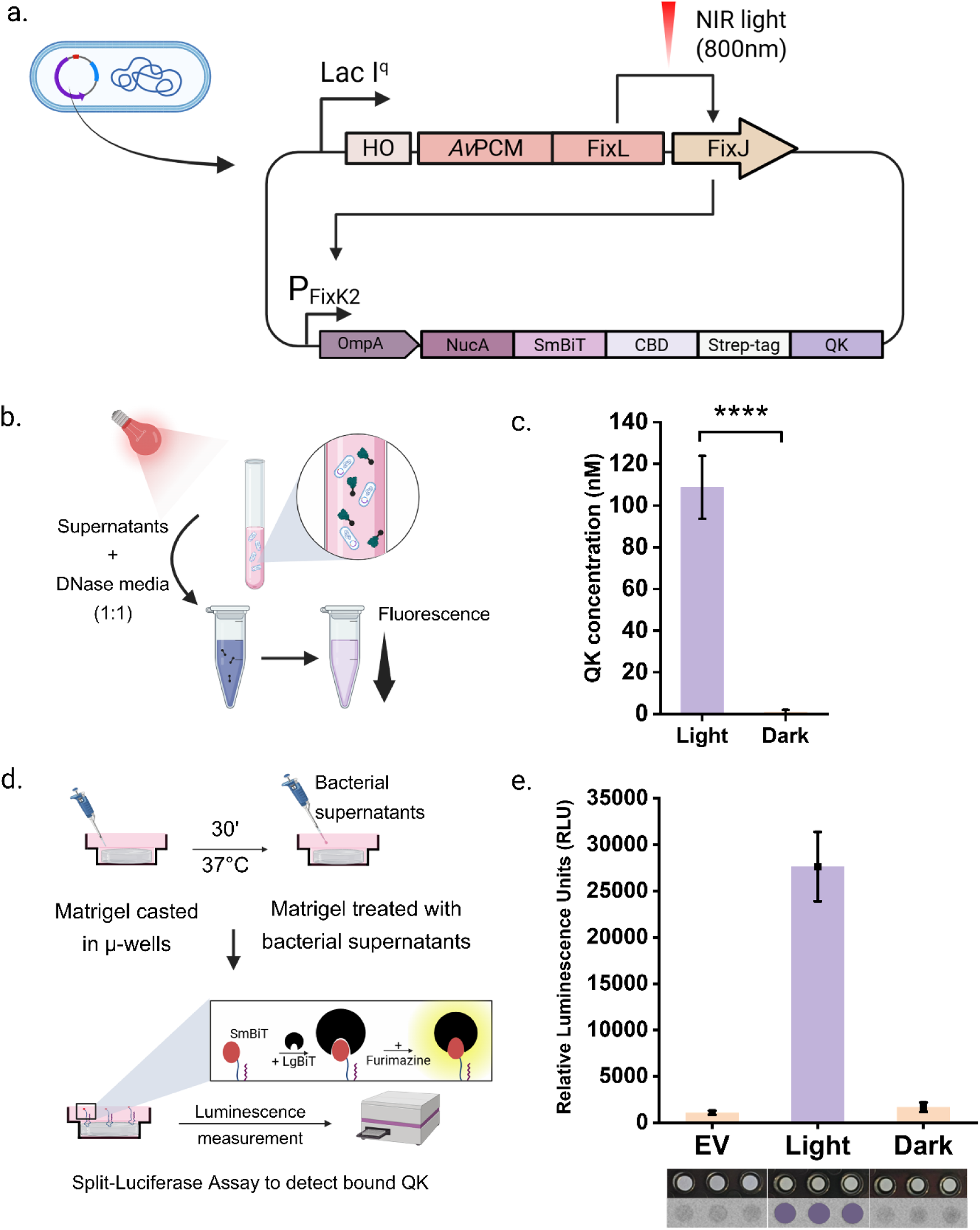
Genetic design and characterization of the near-infrared light responsive bacteria. (a) Architecture of the *Av*NIRusk plasmid for NIR-light-dependent secretion in bacteria. The engineered light-switchable two-component system is constitutively expressed as a tricistronic operon from the lacI^q^ promoter. Said operon contains in sequential order a heme oxygenase for provision of the biliverdin chromophore; a chimeric histidine kinase comprising the photosensor of the bacterial phytochrome from *Agrobacterium vitis* (*Av*PCM) linked to the catalytic domains of *B. japonicum* FixL; and the cognate response regulator FixJ from *B. japonicum*. In NIR light, FixJ is phosphorylated, binds to the FixK2 promoter, and thereby initiates the expression of the therapeutic QK-Fusion protein. Said protein contains the OmpA signal peptide for secretion, the nuclease A from *Staphylococcus aureus* and SmBiT tag for enzymatic and luminescent measurements, respectively. The collagen-binding domain (CBD) enables binding to the extracellular matrix and the QK peptide is a VEGF-mimic. (b) Schematic of Nuclease Assay using culture supernatants. Supernatants from light and dark bacterial cultures were mixed 1:1 with DNase media. Upon incubation, QK-Fusion proteins containing NucA cleaved the DNA causing a loss of fluorescence comparable to the enzymatic activity. (c) Quantification of secreted QK protein under NIR light vs dark conditions. Nuclease assay confirmed strong light-dependent secretion from engineered bacteria (QK concentration in nM), with minimal background under dark conditions. Column height and whiskers represent mean ± SD (N, n=3). (d) Schematic of collagen binding assay. Growth-factor-reduced Matrigel was cast in μ-slides and incubated with bacterial supernatants for QK-fusion to bind to the collagen via the collagen binding domain. Binding was detected via Split-Luciferase Assay. (e) Measurement of Collagen-bound QK using split-luciferase assay. High luminescence signals were detected only in light-induced samples, indicating efficient secretion and collagen binding of QK. Empty vector (EV) and dark conditions showed negligible signal. Column height and whiskers show mean ± SD (N, n=3). (Bottom) Luminescence images of EV, light and dark conditions on a 96-well plate taken with a ChemiDoc^TM^ MP imaging system.

Building upon this platform, we replaced the original reporter protein with a 28 kDa therapeutic fusion protein containing the VEGF-mimetic QK peptide (13 amino acids). This fusion protein also harbors the signal peptide of the native OmpA protein, called *ompa,* to facilitate secretion, a thermostable nuclease from *Staphylococcus aureus*, termed as nuclease A (NucA) domain for enzymatic assays[^25^], an SmBiT tag (11 amino acids) for luminescent quantification [^26^], and a collagen-binding domain (CBD; 10 amino acids) for the protein to bind to the extracellular matrix [^27^] (Figure 1a). This fusion protein is hereafter referred to as QK-Fusion.

Although *E.coli* is a preferred host for recombinant protein production, it exhibits limited efficiency in secreting heterologous proteins due to constraints in the native export systems (^28^,^29^). Particularly in EcN, high secretion can lead to periplasmic burden and growth trade-offs (^30^). While recent efforts have explored signal-peptide optimization and heterologous secretion systems to improve extracellular release, secretion of fully functional recombinant proteins from EcN remains modest. To address this bottleneck, the NucA domain was incorporated not only as a reporter for quantification via DNase assays but also to increase the hydrophilicity of the fusion protein, which improved secretion efficiency by roughly 460 folds (Figure S2a).

Following construction of the *Av*NIRusk-QK-Fusion plasmid and transforming it into EcN to create the EcN NIR-QK-Fusion strain, we evaluated the ability of the engineered bacteria to produce and secrete the fusion protein in a NIR light-responsive manner. To this end, log-phase cultures of these bacteria were incubated either under 800 nm NIR light or dark conditions at 37°C overnight. Culture supernatants were collected, and the quantity of secreted proteins was determined by measuring enzyme activity of NucA through its ability to degrade DNA stained with a fluorescent dye (**Figure 1b**). The NucA activity was quantified by measuring the fluorescence intensity in solution, and the normalized values were converted to concentration values (nM) with a standard curve (Figure S2b). Supernatants from NIR-induced cultures exhibited 137-fold higher DNase activity than those kept in the dark, confirming that the engineered construct was both successfully expressed and secreted in a NIR light-dependent manner (**Figure 1c**). For visual detection of the enzyme activity, we also put the EcN on DNase agar plates containing an indicator dye and kept them either in dark or under NIR light. The plates under NIR light showed a halo of discoloration around the bacteria, confirming that the secreted peptide is capable of degrading DNA. The halo gets bigger by increasing the intensity of incident NIR light (Figure S3).

As stated earlier, activity of the QK peptide relies on its immobilization to the extracellular matrix and hence, we assessed the binding capability of the secreted fusion protein. To test this, culture supernatants collected from EcN NIR-QK-Fusion grown under NIR light or dark conditions were incubated on Matrigel-coated wells and, after washing, immobilization of the protein was visualized via the split luciferase assay. A ChemiDoc^TM^ MP imaging system was used to image luminescence in the wells, and the intensity was quantified using a microplate reader (Figure 1d). Supernatants derived from light-induced cultures showed robust luminescence signals, both visually and quantitatively, in contrast to cultures grown in the dark and negative control. These results confirm that the secreted engineered protein can bind to the matrix in a light-dependent manner (Figure 1e).

### 2.2. ELM Fabrication and Characterization

To translate the genetically engineered bacterial system into a functional and biocompatible ELM, the EcN NIR-QK-Fusion strain was encapsulated within an alginate hydrogel in a core-shell format. The bead architecture closely follows the “Protein Eluting Alginate with Recombinant Lactobacilli” (PEARL) platform which exhibited containment of recombinant Lactobacilli and controlled the release of secreted proteins but failed to contain *E. coli* [^23^]. Thus, modifications to the encapsulation method were developed to ensure containment of EcN.

Encapsulation was carried out in a two-step gelation process. A bacterial-alginate mixture was dropped into calcium chloride to form the core beads, with Alcian Blue added for visualization. These beads were then individually immersed in a second alginate solution and once again crosslinked in calcium chloride to form a uniform shell around the core (Figure 2a). In the macroscopic images, the bead core appears as a uniform blue sphere, while the core plus shell bead is visibly larger, with a transparent outer layer surrounding the core. The corresponding microscopy images give a general view of how the bead looks after 2 days of bacterial growth in the bead (Figure 2b). The bacteria grew into a dense collection of colonies within the core, while not penetrating the shell. Regarding light-responsiveness, fluorescent colonies were observed only in the beads that were irradiated with NIR light (Figure 2c).

**Figure 2.**
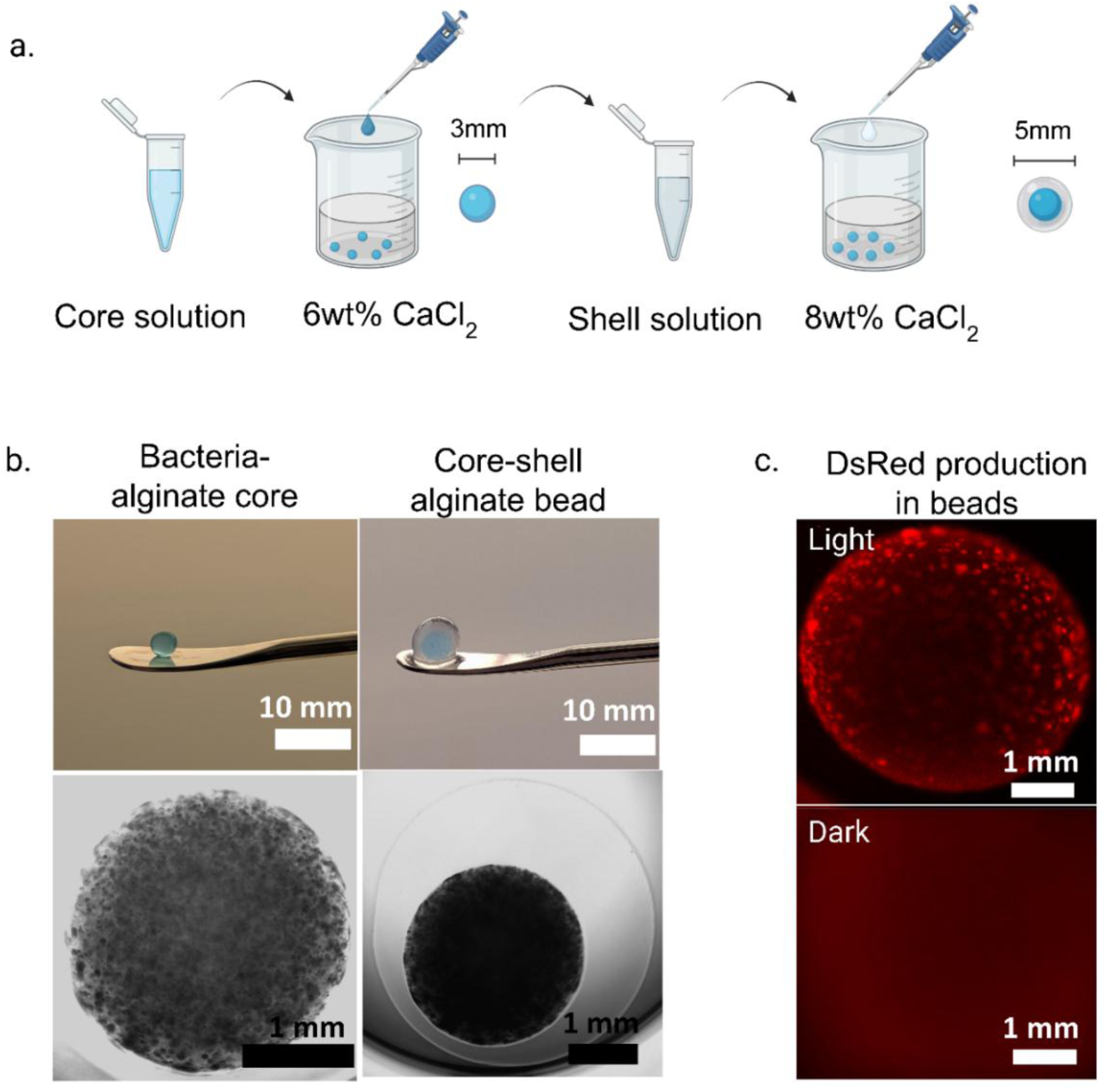
Encapsulation and characterization of engineered EcN in core-shell alginate beads. a. Schematic of the two-step fabrication process of the alginate beads. A bacterial–alginate mixture (1:2) was first crosslinked in 6 wt% CaCl₂ to form the core, followed by a second encapsulation in 1.5 wt% alginate and crosslinking in 8 wt% CaCl₂ to form the shell. b. Macroscopic and brightfield microscopy images showing bead morphology. Top: Macroscopic images of a bacteria-loaded alginate core (left) and a complete core-shell bead (right) taken immediately after crosslinking. Bottom: Brightfield microscopy pictures showing bacterial growth inside the core (left) and core-shell (right) after 24 hours of incubation. c. Fluorescence microscopy of encapsulated EcN expressing *Ds*Red Express2 under NIR light (top) and dark (bottom), confirming light-dependent protein production within the bead.

To reach a formulation that balanced protein release with containment of *E. coli*, we first replicated the published PEARL parameters using EcN in LB medium but observed considerable amount of bacterial leakage within 16 hours of encapsulation. While the PEARL platform evidently sufficed for *Lactobacillus*, it could not contain the fast growing and highly motile EcN. Thus, we systematically optimized the system to achieve containment of EcN. We varied one key parameter at a time such as CaCl_2_ concentration, alginate concentration, crosslinking duration, media composition or bacterial growth phase prior to encapsulation over multiple experimental sets. Table S1 and Figure S4 summarize the empirical workflow that guided the rational formulation of the final core-shell system used in this study. Increasing crosslinking duration to 4 hours and switching from LB to M199 media in the core reduced bacterial escape significantly while supporting secretion. With the escape problem under control, we focused on maximizing protein secretion. We noticed that on increasing alginate concentration, protein secretion decreased further. We hypothesized that the positively charged QK-Fusion might be getting trapped in the negatively charged alginate matrix under higher alginate concentrations or suboptimal crosslinking conditions. Guided by this, we adjusted to lower alginate concentration in the core (2 wt%) and shell (1.5 wt%) and increased CaCl_2_ for shell crosslinking (8 wt%) which improved both protein release and matrix stability. Although no single factor ensured containment on its own, all adjustments together led to the final core-shell system that was stable, reproducible and well-suited for use as an ELM platform. The final formulation consisted of a 2 wt% alginate core (bacteria in M199 media supplemented with 0.04% w/v glucose, OD₆₀₀ = 0.3–0.5) surrounded by a 1.5 wt% alginate shell, with the core crosslinked in 6 wt% CaCl₂ for 30 min and the shell in 8 wt% CaCl₂ for 4–5 h. This optimized system reproducibly generated ≈4 mm beads that supported bacterial viability while preventing bacterial escape for at least 10 days and enabled reliable protein release (Figure 3).

**Figure 3.**
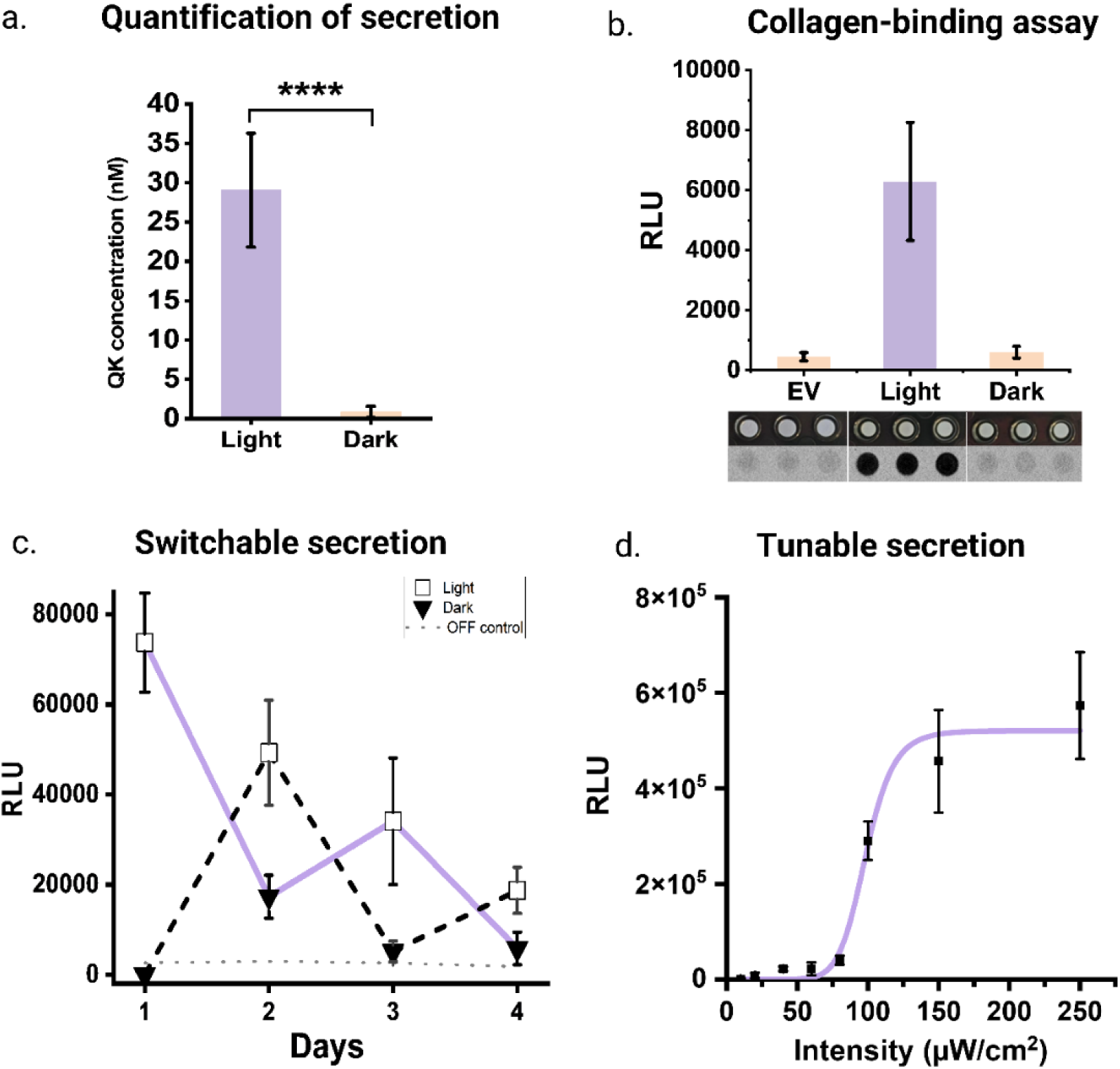
Secretion dynamics and light-responsive control of ELMs. (a) Quantification of secreted QK-Fusion protein from ELMs under light and dark conditions with the Nuclease Assay. (b) Measurement of QK-Fusion bound to the collagen matrix via Split-Luciferase Assay. Luminescence signals were detected only in light-induced bead supernatants, indicating efficient secretion and collagen binding of QK. Empty vector (EV) and dark conditions showed negligible signal. (Bottom) Luminescence images of EV, light and dark conditions on a 96-well plate taken with a ChemiDoc^TM^ MP imaging system. (c) Switchability of QK-Fusion secretion from ELMs across four days under alternating light and dark conditions. Cultures were induced with NIR light (ON) or kept in dark (OFF), and luminescence was measured from supernatants each day. A consistent light–dark response was observed, confirming switchable secretion. QK-Fusion secreting beads kept in dark (OFF) for four days was used as OFF control. (d) Intensity-dependent tunability of QK-Fusion secretion. QK-Fusion secreting beads were subjected to different intensities of 800 nm NIR light. Data were fitted to the Hill equation, yielding *RLU*_*max*_ = 5.2 × 10^5^, *I*_50_ = 98.5 µ*W* ⋅ *cm*^−2^, and *n* = 9.98. For all the graphs, the column height or symbol and whiskers shows mean ± SD, (N, n=3).

### 2.3. Dynamic Light-Controlled Secretion from ELMs

With the encapsulation method established, we proceeded to assess if the system remained inducible and capable of secretion. First, we encapsulated *E. coli* Nissle 1917 harboring an *Av*NIRusk vector driving *Ds*Red expression and exposed some of the resultant beads to NIR light while keeping the remainder in dark. The beads exposed to NIR light showed distinct red fluorescent colonies under the microscope, while those kept in the dark did not (Figure 2c). This confirmed that the bacteria remained alive inside the hydrogel and were viable enough to respond to NIR light and produce the target protein. After demonstrating viability and light-responsiveness, we evaluated whether the system was able to produce and secrete the QK-Fusion protein. Secretion of the protein from the beads was quantified via NucA assay with bead supernatants. After 24 hours of induction, the concentration of the secreted protein was 30 ± 7 nM. In contrast, surrounding medium from uninduced beads (incubated in dark) displayed negligible NucA activity (Figure 3a).

We tested the same supernatants in a collagen-binding assay to evaluate whether the secreted peptide retained its ability to bind to collagen. To this end, we collected the surrounding media from induced, uninduced and control beads, incubated them on Matrigel, and confirmed binding via Split-Luciferase Assay. Supernatants from NIR-induced beads showed strong luminescence signals whereas the uninduced ones showed minimal binding, comparable to the empty vector control. These results indicate that QK-Fusion secreted by the encapsulated bacteria remained intact and functional, retaining its collagen-binding capability after production and release into the surrounding medium (Figure 3b).

To further evaluate the dynamic control of these ELMs, we next tested if our system could reversibly turn secretion ON and OFF over multiple induction cycles. The ELMs were divided into two groups: one exposed to 800 nm NIR light and the other kept in the dark. After 24 hours, the amount of QK secreted in each group was detected using the Split-Luciferase assay. The surrounding media was discarded and replaced with fresh media and the groups of beads were switched - the bead group exposed to light was moved to the dark and vice versa. This cycle was repeated daily for four days. A third group of ELMs, kept in the dark throughout the experiment, served as the continuous OFF control. The experiment showed clear and repeatable switching of QK-Fusion secretion. Beads exposed to NIR light showed sharp increase of luminescence, reflecting active production and release of the QK fusion protein. When moved back to the dark, the luminescence values reached baseline levels, confirming that there was no secretion in the dark. Switching the beads from dark to light once again caused a strong rise in signal. This alternating behavior continued over all four cycles, demonstrating that the secretion response was switchable. Nevertheless, we observed a decrease in ON-state secretion levels as the days progressed, suggesting a possible decline in bacterial viability or activity within the alginate beads under continuous culture in suboptimal M199 medium. Throughout the experiment, the OFF-control group stayed at very low levels, confirming that secretion occurred only when triggered by NIR light (Figure 3c).

We next tested if the secretion from the ELMs was tunable with increasing intensity of NIR light. To confirm this, we exposed the ELMs to a range of intensities from 0 to 250 µW cm⁻². The secretion stayed negligible at low intensities but showed a steep increase beyond 80 µW cm⁻² and plateaued around 150 µW cm⁻² (Figure 3d). This response pattern resembles the previously reported dose response for NIR-responsive EcN producing *Ds*Red [^24^]. In that study, activation was triggered at somewhat lower light intensities, beginning around 20 µW cm⁻² and reaching saturation near 50 µW cm⁻². The present shift to higher intensities for activation might be caused by a combination of factors including decreased penetration of light through dense bacterial colonies in the alginate beads compared to planktonic bacterial cultures, altered metabolic state of the confined bacteria, and slower diffusion of the secreted QK-Fusion protein through the alginate to the surrounding media. However, despite the lowered sensitivity, the light doses required for full system activation remain orders of magnitude below the maximum NIR-light intensity considered therapeutically safe. In addition, lower sensitivity might even prove beneficial in therapeutic applications as it can mitigate unintended activation by low levels of ambient light. Thus, we established that our ELMs can be switched ON and OFF at will and the secretion can be tuned by adjusting the light intensity.

### 2.4. Cytocompatibility of ELM supernatants

Encouraged by the promising results, we proceeded to test the ability of the secreted QK-Fusion protein to drive angiogenic differentiation in human Umbilical Vein Endothelial Cells (HUVECS). Notably, HUVECs are well-established *in vitro* model cells for assessing endothelial cells behaviors such as migration, proliferation and capillary network formation [^31^]. Before performing functional assays, we first examined whether the secreted QK-Fusion protein and other bacterial metabolites were toxic to the HUVECs. We incubated monolayer HUVEC culture for 24 hours with both induced and uninduced culture and ELM supernatants, with supernatants from the empty vector strain-containing ELMs as a control. Cells incubated with the cell culture medium were used as negative control. Two complementary assays were used to assess cytocompatibility. First, we measured cytotoxicity with the lactate dehydrogenase (LDH) assay. A lysis control, in which cells were damaged by Triton X-100 treatment, was used as positive control to estimate maximum cell damage. Second, we measured cell viability with the Alamar Blue assay.

Across all test conditions, cell viability was about 80%, closer to the negative control and far higher than the lysis condition (Figure S5a) Similarly, Alamar Blue assay showed no significant differences between treated and untreated cells. (Figure S5b) These findings confirm that neither QK nor the EcN metabolic byproducts have any adverse effect on HUVECs, and thus, the ELM supernatants are cytocompatible.

### 2.5. Network Formation Assay

Having ruled out cytotoxic effects, we next tested whether the secreted QK-Fusion could drive network formation in HUVEC cell cultures via a 2D network formation assay[^31^]. To this end, we used growth-factor reduced Matrigel in angiogenesis well plates to form a basement membrane-like layer. On the polymerized Matrigel, supernatants from bacterial culture and ELMs were pipetted and incubated at 37°C overnight for the QK-Fusion to bind to the matrix. The following day, serum-starved HUVECs were seeded onto the gels, where they adhered and rearranged to form network-like structures. The network formation was then analyzed by brightfield microscopy after 4-6 hours (Figure 4a).

**Figure 4:**
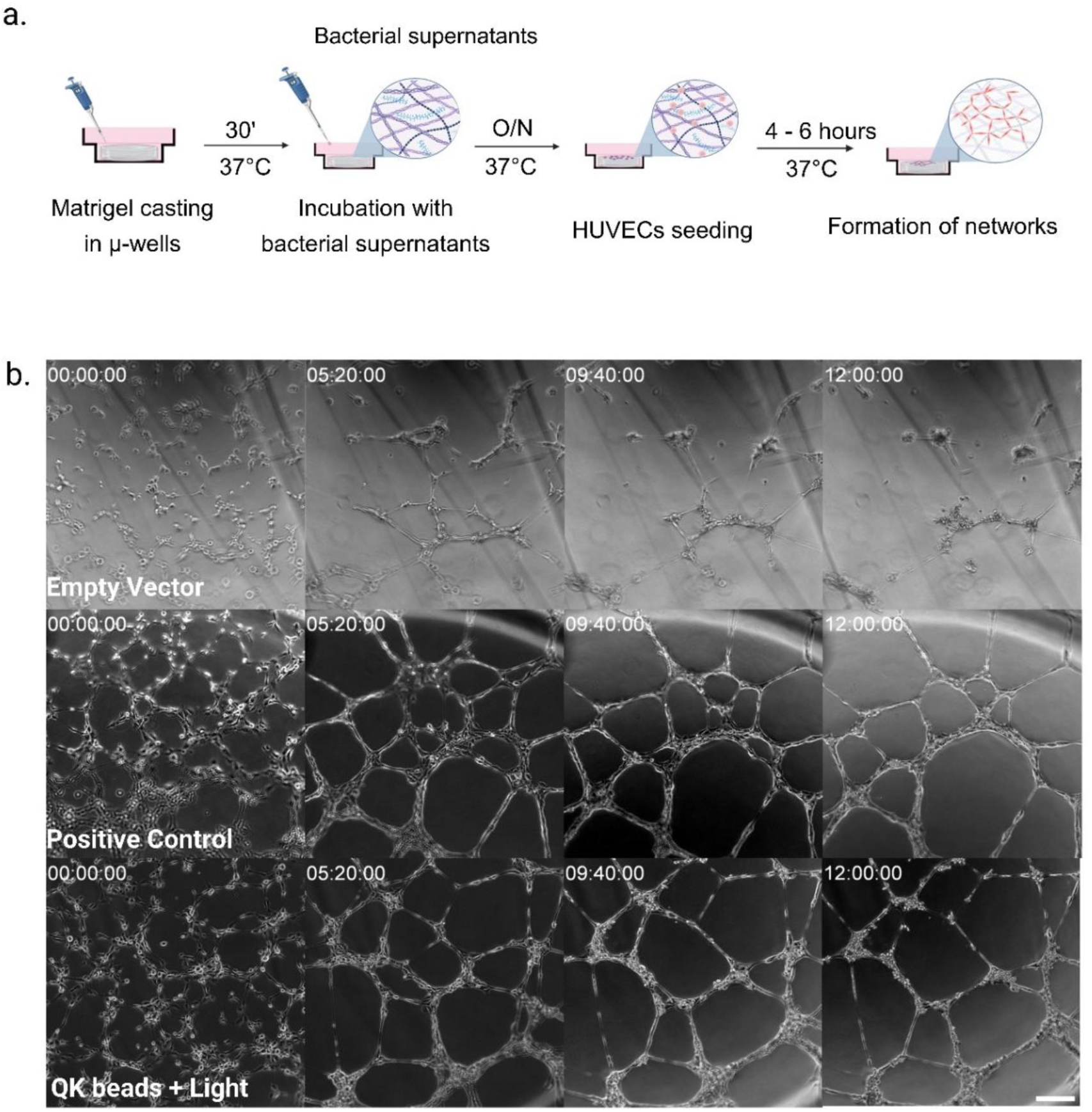
Time-lapse visualization of network formation in HUVECs. (a) Schematic of Matrigel assay performed to analyze 2D HUVEC networks. Growth Factor reduced Matrigel was cast in μ-wells and polymerized for 30 mins, followed by overnight incubation of supernatants from cultures and ELMs in different conditions. The next day, HUVECs were seeded onto the Matrigel and network formation was tracked for 4-6 h. (b) Representative time-lapse brightfield snapshots showing network formation in three experimental conditions: negative control (M199 media, top), positive control (M199 media +supplements + FBS, middle) and supernatants from light-induced ELMs (bottom) at timepoints 00:00, 05:20, 09:40 and 12:00 (hh:mm). Scale bar denotes 200 µm.

As this is a highly dynamic process, we first tracked HUVEC arrangement over time by time-lapse brightfield microscopy (Figure 4b). Cells treated with supernatants from negative control, positive control, and NIR-induced QK beads were followed over 20 hours while capturing still images every 20 minutes. Representative snapshots from selected timepoints (00:00, 05:20, 09:40 and 12:00 in hh:mm) are shown in Figure 4b. The full videos are shown in Supporting Information (Movies S1-S3). The positive control condition triggered a rapid rearrangement of the endothelial cells into well-defined networks with big lumens which continued to mature over time. The networks formed in this condition were stable over 12 hours. On the other hand, the negative control condition showed little to no organization. Small groups of cells reorganized in an unstable manner, which still shows the suitability of Matrigel to host endothelial cells even in the growth factor depleted version. Interestingly, HUVECs treated with supernatants from light-induced ELMs formed well-defined networks that closely resembled the networks formed by positive controls in terms of morphology and stability over time (stable up to 7 hours).

From the time-lapse results we established that network formation occurs during the first 4-6 hours (Figure 5). Building upon this, we extended our analysis to incorporate multiple conditions and assess network morphology via endpoint microscopy. We compared the potential pro-angiogenic effects of the secreted QK-Fusion from culture and ELM bead supernatants in light and dark conditions, together with the empty vector and positive and negative controls. The microscopy images revealed distinct morphological differences between the experimental groups (Figure 5a). The positive control presented elongated networks, with connected and thick vessel-like structures, while the negative control showed scattered cells without proper connected vessels. In contrast, supernatants from both culture and ELMs incubated under NIR light supported robust network formation closely resembling the positive control. In comparison, supernatants from culture and ELMs in the dark induced only limited networks — more than the negative control but much lower than the light-induced conditions. This effect is likely due to low levels of leaky expression in non-induced conditions.

**Figure 5.**
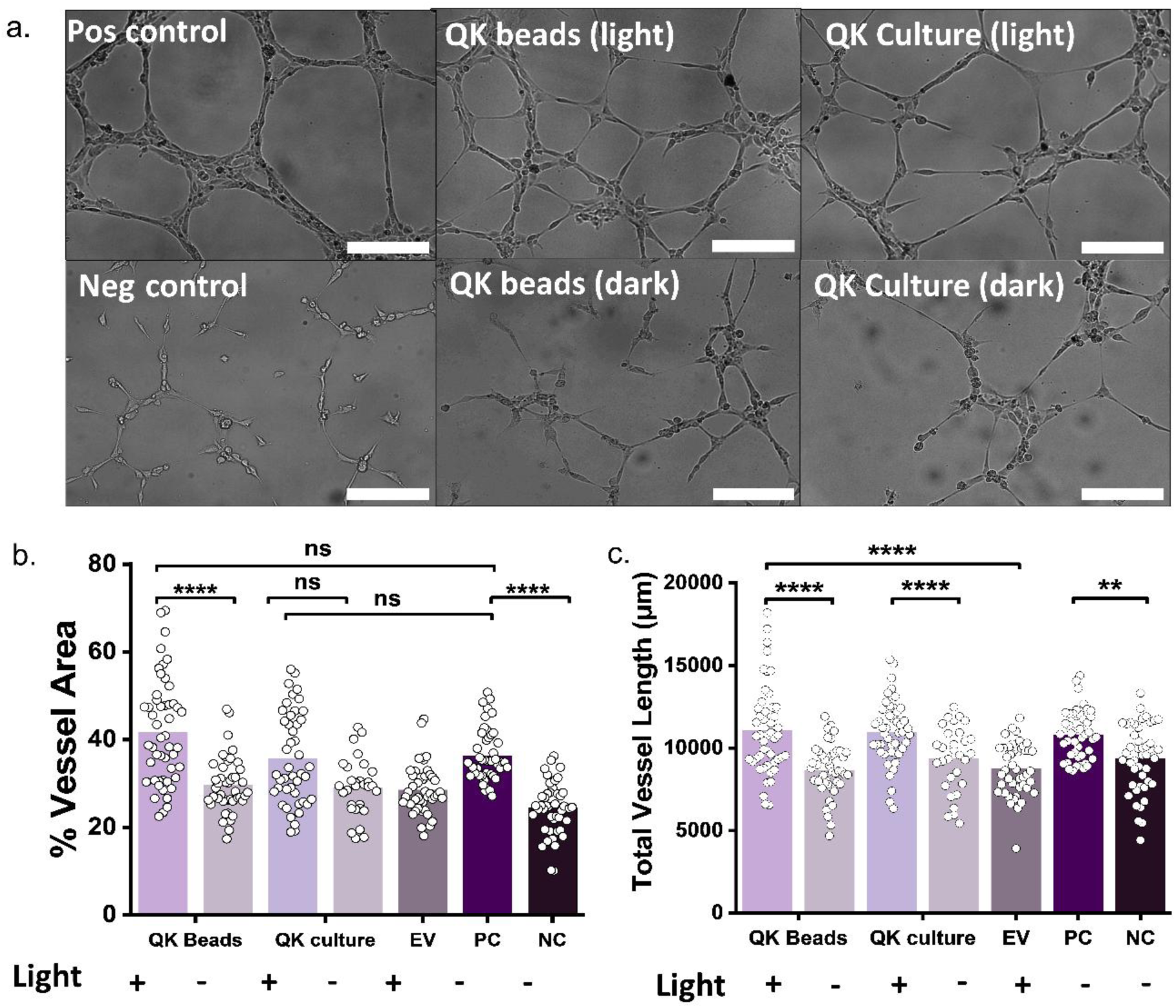
QK-Fusion mediated HUVEC network formation from ELM and culture supernatants. (a) Representative brightfield images showing network morphology after treatment with different supernatants. Light-induced ELM and culture supernatants promote longer, continuous network formation similar to the positive control whereas supernatants from ELMs and culture kept in the dark form shorter and discontinuous networks like the negative control. Brightness and contrast of the images have been adjusted for maximal visualization. Scale bar denotes 200 μm (b-e) Quantitative analysis of angiogenic parameters using ImageJ and AngioTool: (b) Percentage of vessel-covered area (c) Total vessel length (μm). Bars represent mean value, dots represent each individual datapoint (45 datapoints, derived from N= 3 biological replicates × n=3 technical replicates × 5 fields of view) (EV: Empty Vector, PC: positive Control, NC: Negative Control)

Quantitative analysis of network formation was performed using Angiotool in ImageJ (Figure 5b-e). We focused on two parameters to effectively assess angiogenic behavior across the different conditions. Vessel Area (in percentage), which measures the fraction of the image occupied by segmented vessels, and Total Vessel Length, defined as the cumulative length of all skeletonized vessel segments. These measures together reflect the coverage of the network and the quality of the vessels [^32^].

The average percentage of area covered by vessels (percentage vessel area) was the highest in light-induced beads, and both light-induced culture and beads were comparable to the positive control, suggesting strong pro-angiogenic effects (Figure 5b). Under dark conditions, both beads and culture showed an average percentage of vessel area above the negative control and empty vector (although not significantly different), which is consistent with the low-level background activity seen in the images (Figure 5a). The average values measured for the total vessel length also reflected the improved vessel structures formed in QK beads and cultures, showing comparable values to the ones measured for the positive control (Figure 5c). In contrast, the dark conditions formed short, discontinuous vessels similar to negative control and empty vector. This confirmed that the secreted QK-Fusion aided in the formation of the longer capillary-like structures.

Taken together, QK-Fusion secreted from the light-induced ELMs favored better network formation over all other conditions even though the amount of peptide secreted is significantly lower than the amount of peptide secreted in culture. This partially aligns with the previously reported findings by Nakatsu *et al*., who showed that lower concentrations of VEGF₁₆₅ lead to thinner, more organized, and stable vessels, while higher concentrations result in wider, disorganized structures [^33^]. In our case, the ELMs seemed to be delivering QK at a more optimal concentration that supported proper vessel growth compared to the higher QK levels present in culture supernatants, which likely saturated the system, leading to less effective network formation.

## 3. Conclusions

Chronic wounds remain a major therapeutic challenge due to persistent inflammation and impaired angiogenesis [^34^]. Near-Infrared photobiomodulation offers a non-invasive means to address these barriers due to its deep tissue penetration, mitochondrial ATP synthesis stimulation and anti-inflammatory effects [^35^]. Complementing NIR therapy with peptidomimetics such as VEGF-derived QK provide stable and cost-effective alternatives to full-length growth factors for controlled vascular repair.

Building on this potential, we developed a NIR-responsive Engineered Living Material utilizing the regenerative benefits of NIR therapy with the precision of synthetic biology. The probiotic *E.coli* Nissle 1917 was engineered with an optogenetic circuit that can secrete the QK-Fusion protein in response to 800 nm NIR light. Encapsulation of the engineered EcN alginate-based core-shell beads ensured bacterial viability, biocontainment and sustained release of therapeutic. The resulting construct demonstrated robust, reversible and intensity-dependent secretion thus achieving both temporal and spatial control. The encapsulated systems outperformed culture-based systems in forming elongated, organized, and interconnected networks of HUVECs, despite secreting lower overall amounts of peptide. Moreover, the encapsulated format reduced background leakiness in the dark state, further supporting its suitability for precision delivery.

Overall, our findings establish a modular, optogenetically controlled ELM platform capable of programmable, spatially confined delivery of pro-angiogenic cues. The use of deep-tissue-penetrant NIR light, clinically approved bacterial strains, and scalable encapsulation methods together point toward translational potential for treating ischemic or non-healing wounds with unprecedented precision and responsiveness. Of decisive advantage, the use of NIR light for photobiomodulation is well established, considered therapeutically safe, and implementable at low cost. Our present work paves the way towards innovative treatment strategies that combine photobiomodulation and NIR-light-responsive, programmable bacteria to achieve added or even synergistic effects.

## 4. Materials and Methods

### Bacterial Strain, Media and Plasmids

#### AvNIRusk- QK-Fusion plasmid creation

The QK-Fusion construct was ordered as an eBlock from Integrated DNA Technologies (IDT) and codon-optimized using the IDT codon Optimization Tool. The gene was cloned into the *Av*NIRusk plasmid (pMH028, ^36^) replacing the *Ds*Red reporter gene using NEBuilder HiFi Assembly cloning kit (New England Biolabs GmbH, Germany, E5520S). For creation of the empty vector (EV) control, the *Av*NIRusk plasmid was amplified excluding the *Ds*Red gene and circularized using the KLD Enzyme Mix (New England Biolabs GmbH, Germany, E0554S). All plasmids were initially transformed into NEB 5-alpha Competent *E*. *coli* cells (New England Biolabs GmbH, Germany, Art. No. C2987) and plated on LB agar supplemented with kanamycin ((kanamycinsulfat, Carl Roth GmbH, Germany)). Positive clones were screened by colony PCR, followed by plasmid extraction (Thermo Fisher Scientific, K210011) and Sanger sequencing (Eurofins Genomics) to verify the sequence of the construct.

#### Preparation of Chemically Competent <u>E. coli</u> Nissle 1917

An overnight culture of wild-type *E. coli* Nissle 1917 grown at 37°C, 250 rpm was subcultured into 50 mL of fresh LB to a starting O.D_600_ of 0.01. The culture was grown under same conditions until log phase (OD₆₀₀ ≈ 0.3-0.4). The cells were then chilled on ice and pelleted by centrifugation at 4000 rpm for 10 min at 4 °C. To prepare the cells for transformation, the pellet was washed twice with 10 mL of freshly-prepared, ice-cold 200 mM calcium chloride solution (Carl Roth GmbH, Germany), followed by a single wash with 10 mL of a 1:1 mixture of 200 mM CaCl₂ and 10% (w/v) glycerol. After the final wash, the pellet was gently resuspended in 1 mL of the same CaCl₂:glycerol mixture. Aliquots of 50 µL were prepared and either used immediately or stored at –80 °C for future transformations.

#### Transformation into <u>E. coli</u> Nissle 1917

Sequence-verified *Av*NIRusk-QK-Fusion plasmid and empty vector plasmids were transformed into EcN via heat-shock. Briefly, about 500 ng of plasmid DNA was gently mixed with 50 µL of chemically competent *E. coli* Nissle 1917 cells and incubated on ice for 30 minutes. The mixture was then subjected to a heat-shock at 42°C for 30 seconds using a water bath, followed by immediate cooling on ice for 5 minutes. 900 µL of pre-warmed SOC medium was added to the cells, and the suspension was incubated at 37 °C for 1 hour with shaking to allow recovery. After recovery, 200 µL was spread onto LB agar plates containing 50 µg/ml kanamycin and the plates were incubated at 37°C overnight. All procedures were performed under dark or low-light conditions to avoid unintended activation.

### Bacterial performance testing

#### Bacterial culture for secretion

EcN carrying the desired plasmid was grown from cryo-culture stocks in Luria-Bertani (LB) medium supplemented with kanamycin (50 µg mL⁻¹) at 37 °C with shaking at 250 rpm for 16 h. The overnight culture was diluted to an initial OD₆₀₀ of 0.01 into fresh LB medium with kanamycin under two conditions: induced and uninduced. Cultures were incubated at 37 °C until mid-log phase (OD₆₀₀ = 0.6–0.8), followed by centrifugation at 4000 rpm for 10 min at room temperature. The resulting cell pellets were either resuspended in LB medium or Medium 199 (M199; Gibco) containing kanamycin (50 µg mL⁻¹).

NIR illumination was performed using a custom-built LED setup consisting of six 800nm LED bulbs (LED800-01AU, Roithner Lasertechnik) arranged in a 3 × 2 matrix on a flat base. The base was fitted with a holder accommodating six 10mL culture tubes positioned at a 30° tilt, maximizing bacterial growth. The LEDs were connected to a control unit containing a master switch and three independent channels, each governing a pair of bulbs. Each channel was equipped with an individual on/off switch and a potentiometer allowing fine-tuning of light intensity across 10 major divisions, with 100 incremental steps per division. The system was calibrated using a power meter to deliver a light intensity of 2 mW cm^-2^ at the sample plane during induction experiments.

For induction, cultures were exposed to near-infrared (NIR) light at 800 nm with an intensity of 2 mW cm⁻² for 20-22 h at 37 °C. Uninduced samples were protected from light using aluminum foil.

#### Fluorescence validation of supernatants on DNase Agar plates

To validate fluorescence of secreted proteins and visually confirm that *Ds*Red expression does not induce DNA degradation, a DNase agar plate supplemented with kanamycin (50 µg/mL) was divided into three zones. Onto each zone, 5 µL of mid-log phase bacterial cultures were spotted: (i) *E. coli* Nissle 1917 expressing QK-Fusion, (ii) EcN expressing *Ds*Red, and (iii) an empty vector (EV) control. The plate was incubated under NIR light to induce expression and imaged after 16–18 h using a ChemiDoc^TM^ MP imaging system (Bio-Rad, Germany).

#### Fluorescence quantification

*E. coli* Nissle 1917 carrying the *Av*NIRusk–*Ds*Red construct was first grown overnight in LB medium at 37 °C with shaking (250 rpm). The next day, cultures were diluted to OD₆₀₀ ≈ 0.01 in fresh LB medium supplemented with kanamycin and incubated until mid-log phase (OD₆₀₀ = 0.4–0.6). The cultures were pelleted down (4000 rpm, 10 mins) and resuspended in same volume M199 medium supplemented with kanamycin. Cultures were then exposed to 800 nm NIR light or kept in the dark overnight at 37 °C. The following day, 500 µL from each culture was harvested, pelleted (4000 rpm, 5 min), and resuspended in PBS to a final OD₆₀₀ of 1.0. A volume of 200 µL of each normalized suspension was transferred into black, clear-bottom 96-well plates (Greiner Bio-One, Germany). Fluorescence was measured on a TECAN microplate reader (Infinite 200 Pro, Tecan Deutschland GmbH, Germany) using excitation at 500 ± 9 nm, emission at 554 ± 9 nm. Fluorescence values were normalized to OD₆₀₀ to account for cell density differences.

#### Light-Dose-Response Assay

To assess the NIR light induced expression of the fluorescent reporter *Ds*Red in *E. coli* Nissle 1917 light-dose-responses were recorded as before [^24^]. In brief, bacteria containing *Av*NIRusk-*Ds*Red or an empty vector control were grown in 5 mL LB medium supplemented with 50 µg mL⁻¹ kanamycin (LB-Kan) at 30°C and 225 rpm agitation for 21-24 h in darkness (non-inducing conditions). All subsequent steps were performed under green safe light. The 100-fold diluted bacteria were distributed in 200 µL per well in a black-walled microtiter plate with clear bottom (µClear, Greiner BioOne, Frickenhausen, Germany). Different NIR light intensities were applied using a custom made 8-by-8 LED matrix reported in [^37^] emitting at (800 ± 18) nm controlled by an Arduino circuit board. The light intensities were calibrated with a power meter (model 842-PE, equipped with a 918D-UV-OD3 silicon photodiode, Newport, Darmstadt, Germany). After an 18 h incubation at 37°C and 750 rpm agitation, the OD_600_ and *Ds*Red fluorescence with excitation and emission wavelengths of (500 ± 9) nm and (554 ± 9) nm were recorded with a Tecan Infinite M200 Pro MTP reader (Tecan Group, Ltd., Männedorf, Switzerland). The *Ds*Red fluorescence was normalized by OD_600_ and the empty vector background was subtracted. Data are represented as a function of the light intensity averaged over the duty cycle and are described by Hill isotherms using the Fit-o-mat software [^38,39^].

#### Nuclease A Secretion Assay on Agar Plates

The Difco^TM^ DNase test agar (Fisher scientific) was prepared according to the manufacturer’s instructions and 0.05 g L⁻¹ of the methyl green indicator (Serva) was added manually. A bacterial colony containing the *Av*NIRusk-NucA was resuspended in 20 µL phosphate-buffered saline and 1 µL was dropped per agar plate. The plates were incubated at 37°C overnight in darkness or under constant NIR light applied with the Arduino-controlled custom-built LED matrix.

#### NucleaseA assay with supernatants

DNase media was prepared by precipitating agar from the commercial DNase Agar (Altmann Analytik GmbH, Germany), followed by sterilization by autoclaving. The final formulation comprised of Tryptose (20 g/L), Sodium Chloride (5 g L^−1^), Calf Thymus DNA (2 g L^−1^) and Methyl Green (0.05 g L^−1^). For measuring secretion from bacterial cultures, the supernatants from the uninduced and induced cultures were filter-sterilized using 0.22µm syringe filters. They were then mixed with DNase media in a 1:1 ratio for a final volume of 800 µl. Following preparation, tubes were wrapped in aluminum foil to prevent light-induced degradation of methyl green. All samples were incubated at 37 °C with shaking at 250 rpm for 4 h. After incubation, 200 µL from each dilution was transferred in triplicate to a black 96-well plate with a transparent bottom. Fluorescence was measured using a Microplate Reader (Infinite 200 Pro, Tecan Deutschland GmbH, Germany) at an excitation/emission wavelength of 633/668 nm, with the Z-position set at 19,000 µm and gain at 100.

#### Standard curve preparation

For the standard curve preparation, DNase media was added to the cell free supernatant of *E. coli* Nissle 1917 harboring the Empty Vector, collected after 16 h of NIR light induction, in a 1:1 ratio. Using this solvent as a base, serial dilutions of pure NucleaseA protein (Sigma-Aldrich) were prepared in 1.5 mL Eppendorf tubes for concentrations of 200, 100, 50, 25, 12.5, 6.25, 3.12, and 1.56 nM, for a final volume of 800ul. After incubation, 200 µL from each dilution was transferred in triplicate to a black 96-well plate with a transparent bottom. Fluorescence was measured using a Microplate Reader (Infinite 200 Pro, Tecan Deutschland GmbH, Germany) at an excitation/emission wavelength of 633/668 nm, with the Z-position set at 19,000 µm and gain at 100.

#### Matrigel preparation

Growth Factor Reduced (GFR) Matrigel® (356231, Corning) aliquots were thawed overnight at 4 °C. A box of 10 µL filter tips was also pre-chilled at 4 °C to prevent premature polymerization during handling. The following day, 10 µL of cold Matrigel solution was carefully dispensed into each well of a pre-chilled 96-well 3D µ-Plate (Ibidi) on ice. The plate was then incubated at 37 °C for 30–45 min to allow complete gelation.

#### Collagen Binding Assay

Supernatants from overnight induced and uninduced bacterial cultures were collected by centrifugation at 4000 rpm for 10 mins. 50 µl of these supernatants were then pipetted in each well and the plate was incubated in the dark at 37°C for 90 minutes. After incubation, each of the wells was washed thrice with sterile PBS. Each wash was performed very carefully to not disrupt the Matrigel surface. Matrigel-bound QK-Fusion was quantified by Split-Luciferase Assay. Empty vector culture supernatant was used as a negative control.

#### Split-Luciferase Assay

Split-luciferase experiments were performed in white, 96-well flat-bottomed microplates and measured using a microplate reader (Infinite 200 Pro, Tecan Deutschland GmbH, Germany). The measurements were performed in triplicates with an integration time of 1000 ms for each measurement. LgBiT protein was always freshly thawed and diluted to 1μM in PBS. The substrate solution was prepared by diluting Furimazine (Nano-Glo® Luciferase Assay Substrate, Promega) in Nano-Glo® Luciferase Assay Buffer in a 1:50 ratio. The substrate solution was always freshly prepared, protected from light and directly added prior to the measurements. In the well plates, 12.5 μL of LgBiT was added to 12.5 μL of each sample supernatant. Consequently, to each of the wells, 25 μL of the substrate solution was added and the luminescence measurements were performed in the plate reader after incubating the plate in the dark for 10 min. The plate was also imaged in the ChemiDoc on the chemiluminescence channel. Empty vector supernatants were used as negative control.

### Fabrication and characterization of ELMs

#### Bacterial Culture for ELMs

Escherichia coli Nissle 1917 (EcN) carrying the desired plasmid was grown from glycerol stocks in Luria-Bertani (LB) medium supplemented with kanamycin (50 µg mL⁻¹) at 22 °C with shaking at 250 rpm for 16 h.

#### Fabrication of ELMs

The core-shell alginate beads were fabricated following the protocol described in [^23^] with certain modifications for encapsulating *E. coli* Nissle 1917. Briefly, *E. coli* Nissle 1917 cultures were grown to mid-log phase (OD₆₀₀ = 0.3–0.5), harvested by centrifugation at 4000 rpm for 10 min, and resuspended in Medium 199 (M199, Gibco) containing 50 µg mL⁻¹ kanamycin and 0.4 w/v% glucose. The bacterial suspension was mixed with 3 wt% alginate in a 1:2 volume ratio to form a 2 wt% alginate–bacteria mixture, supplemented with 1% v/v Alcian blue for visual tracking. Droplets (10 µL) were dispensed into a 6 wt% CaCl₂ solution and allowed to crosslink for 30–45 min, forming the alginate core beads.

To generate the shell layer, a 1.5 wt% alginate shell solution was prepared by diluting the 3 wt% alginate stock 1:1 with Milli-Q water and vortexed for uniformity. Each previously formed alginate core bead was dipped into the shell solution and transferred—along with approximately 25 µL of shell solution—into an 8 wt% CaCl₂ crosslinking bath using a modified pipette tip (tip cut to widen the bore). The core–shell beads were allowed to solidify in CaCl₂ for 4-5 h at room temperature to ensure complete ionic crosslinking. Finally, beads were washed with sterile Milli-Q water and transferred into a 96-well plate with 250 uL of M199 media + kanamycin in each well. The plate was covered in aluminum foil and left overnight in a 37°C incubator for the bacteria to grow.

#### Switchable secretion

Core–shell alginate beads containing *E. coli* Nissle 1917 engineered for NIR light-responsive QK-Fusion expression were incubated overnight at 30 °C in M199 medium supplemented with 50 µg mL⁻¹ kanamycin. Blank beads devoid of any bacteria and beads with empty vector containing *E. coli* Nissle 1917 were used as controls. The following day, beads were distributed into two sterile 96-well plates. Plates were incubated under either 800 nm NIR light at 2 mW cm⁻² or complete darkness (wrapped in aluminum foil) for 24 h. Following incubation, 25 µL of cell-free supernatants were collected from each well for analysis. The remaining medium was completely replaced with fresh supplemented M199, and the light conditions were switched: the plate previously exposed to light was transferred to dark conditions and the plate previously kept in the dark was moved to the NIR light source. This light/dark switching cycle was repeated every 24 h for four consecutive days, with supernatant collected daily. The peptide detection was done with the help of Split nanoluciferase assay by adding 25 μL of LgBiT and 50 μL of diluted substrate to the 25 μL of supernatant and measuring luminescence signals in the plate reader after 10 minutes of incubation in the dark.

#### Light Dose-response assay with ELM supernatants

To characterize the dose-responsive behavior of our NIR-controlled system, we measured QK secretion under a gradient of light intensities (0, 10, 20, 40, 60, 80, 100, 150, 250 μW·cm⁻²), analogous to the approach described earlier. Fresh beads were allowed to grow for two days at 30°C. They were then induced under each defined intensity for 24 h using a custom made 8-by-8 LED matrix [^37^] emitting at (800 ± 18) nm controlled by an Arduino circuit board. Following induction, supernatants were collected, and 25 µL of each were assayed by split-luciferase to quantify secreted QK-Fusion. The resulting intensity–response data were fitted in Origin v9.9 using a Hill model (*y* = *V*_*max*_ *X*^*n*^⁄(*k*^*n*^ + *X*^*n*^)), yielding *RLU*_*max*_ = 5.2 × 10^5^, *I*_50_ = 98.5µ*W* ⋅ *cm*^−2^, where *I_50_* refers to the light dose required for half-maximal activation, *n* = 9.98, and *R*^2^ = 0.93, confirming the tunable, sigmoidal activation of secretion with increasing NIR intensity.

### HUVECs culture and network formation

#### HUVECs culture conditions

Human umbilical vein endothelial cells (pooled donors, HUVECs; Lonza) were cultured on gelatin-coated flasks (0.2% w/V) to enhance adherence. Cells were maintained in Medium 199 supplemented with 20% fetal bovine serum (FBS gold; PAN-Biotech, Germany), 1% penicillin-streptomycin (Sigma), 12 μg/mL endothelial cell growth supplement (ECGS; Sigma, C-30160), and 0.1 mg/ml sodium heparin (Sigma, H-3149).

#### HUVECs passaging and seeding

For routine passaging, spent media was discarded from the culture flask and washed thoroughly with sterile PBS solution. For detachment of cells, approximately 2 mL/10 cm^2^ pre-warmed trypsin was added to the flask and incubated at 37°C for 1-2 min till 90% of the cells were visibly detached. To stop enzymatic activity, pre-warmed complete growth medium was added at twice the volume of trypsin used. The cells were then collected in a sterile falcon and centrifuged at 1000 rpm for 7 min. After centrifugation, the supernatant was discarded, and the cell pellet was resuspended in 1 mL of complete growth medium. For counting cells, 25 μL of the cell suspension was mixed with 25 μL of Trypan Blue dye (Thermo Fisher Scientific). 10 μL of this mixture was then loaded onto the cell counter (CellDrop) for counting cell number and cell viability. If the cell viability was >75%, the cells were seeded at a density of 1000 cells per cm^2^ in complete growth medium in flasks lined with 0.2% w/v of gelatin. Cells were then incubated at 37°C in a humidified atmosphere containing 5% CO₂, and medium was changed every 48 h or as required.

#### Cytocompatibility assays

to prepare for evaluating cytocompatibility, HUVECs (5000cells/well) were seeded in a sterile 96-well μ-plate (Ibidi, Cat.No: 89646) in 50 μL growth media and allowed to attach and form a monolayer (37°C, 20 h, 5% CO_2_). The spent media was then removed and treated with bacterial supernatants from different conditions (light and dark-treated bacterial culture, light and dark-treated beads and empty vector beads). The cells were incubated overnight at 37°C and 5% CO_2_.

#### Cytotoxicity Assay

Cytotoxicity was assessed via a Lactate Dehydrogenase Assay (LDH) using the CytoTox 96® Non-Radioactive Cytotoxicity Assay (Promega). according to the manufacturer’s protocol. In brief, 30 µL of supernatant from each experimental well was transferred to a fresh 96-well plate and mixed with an equal volume (30 µL) of LDH substrate solution. The reaction was allowed to proceed for 20 min at room temperature in the dark, after which 30 µL of stop solution was added to terminate the reaction. Absorbance was measured at 490 nm using a TECAN Spark microplate reader. Cells cultured in standard growth medium served as the negative control, while cells lysed with 37.5% Triton X-100 were used as the positive control. Media-only wells were included to correct for background absorbance, termed as ‘blank’. These blank values were subtracted from all the sample and control values. The percentage of cell death was then calculated as:

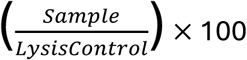

#### Cell-viability Assay

Cell viability was assessed using the AlamarBlue™ reagent (Invitrogen) according to the manufacturer’s instructions. In brief, a 10% (v/v) solution of AlamarBlue in growth medium was added to cells treated with the supernatants. Cells maintained in standard growth medium served as positive control, while medium-only wells were included as blanks. Fluorescence (Ex/Em 570/600 nm) values were recorded using a TECAN Spark microplate reader. Blank values were subtracted from all the sample and control values.

#### Network Formation Assay

For the endothelial network formation assay, growth factor-reduced Matrigel (Corning) was thawed overnight at 4 °C. A volume of 10 µL of Matrigel was dispensed into each well of angiogenesis assay plates on ice and incubated at 37 °C for 30–45 minutes to allow polymerization. Following gelation, 50 µL of treatment solution was added to each well in triplicate and incubated overnight at 37 °C. HUVECs at 60–70% confluency was cultured in Medium 199 (M199) supplemented with 5% fetal bovine serum (FBS) and 1% penicillin-streptomycin (P/S) for 20 h. After this conditioning period, cells were harvested by trypsinization and were neutralized using pre-warmed complete medium. Cells were pelleted by centrifugation and resuspended in M199 supplemented with 1% P/S. The suspension was adjusted to a final density of 15,000 cells in 50 µL. Prior to cell seeding, the treatment-containing medium was removed from the Matrigel-coated wells. The cell suspension (50 µL) was gently added to each well and allowed to attach for 1 h at 37 °C. Plates were then incubated at 37 °C in a humidified 5% CO₂ incubator for 6 h, and network formation was monitored hourly using bright-field microscopy.

#### Image Acquisition for timelapse imaging

To monitor HUVEC network formation in real time, live-cell imaging was conducted using an Observer Z1 inverted wide-field microscope (Carl Zeiss, Oberkochen, Germany), equipped with a temperature-controlled incubation unit maintained at 37 °C and a 5% CO₂ atmosphere. The µ-slides were prepared as previously described and immediately transferred to the microscope’s incubation chamber. Phase-contrast images were captured using a Zeiss EC Plan-Neofluar 10×/0.3 Ph1 objective lens, without additional magnification, at 20-minute intervals over 22 hours. Image acquisition was performed using a Prime BSI Express sCMOS camera (Teledyne Photometrics, USA), with ZEN software (version 3.5, ZEISS) controlling the imaging process. For each experimental condition, three non-overlapping fields per well were imaged, with a minimum of two biological replicates.

#### Image Acquisition for endpoint microscopy

Brightfield images from the Matrigel-based network formation assay were acquired using Leica DMI6000 B epifluorescence microscope at 10X magnification after 4hours. Five non-overlapping fields were imaged per well, with at least three biological replicates per condition. The experiment was independently repeated three times across all seven experimental groups.

#### Image analysis

Brightfield images from the Matrigel-based angiogenesis assay were acquired using Leica DMI6000 B epifluorescence microscope at 10X magnification. Five non-overlapping fields were imaged per well, with at least three biological replicates per condition. The experiment was independently repeated three times across all seven experimental groups. Image analysis was performed in Fiji (ImageJ) 2.16.0/1.54p using the Trainable Weka Segmentation (TWS) plugin and the AngioTool plugin. The AngioTool plugin extracted the following parameters: vessel area, total vessel length, junction density and number of endpoints and junctions. Vessel area was calculated as the fraction of the image occupied by vessel-like structures, Total vessel length was calculated as the sum of all detected vessel segment lengths, junction density was calculated as the number of branching points per unit image area and the number of endpoints was calculated as per Equation 1. A custom-written macro (provided in the Supplementary Information) enabled batch processing: it applied a pre-trained Weka model for vessel segmentation, thresholded the images, and generated binary masks for downstream analysis.

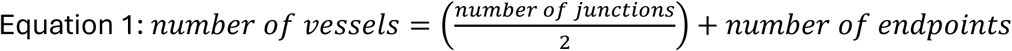

### Statistical Analysis

Statistical analysis was carried out using GraphPad Prism v9 software and plotted using Origin 2022 v9.9 software. All experiments were repeated independently three times (N=3 biological replicates) unless otherwise noted. Samples were assayed in triplicates (n=3 technical replicates) unless otherwise noted. All graphs represent mean ± standard deviation (SD) unless otherwise noted. Students’ T-tests were used to determine significant differences between the means of the groups. For the network formation analyses, the goodness of fit of the data was tested via Shapiro-Wilk normality test. When comparing three or more groups: normal distributed populations were analyzed via analysis of variance test (ANOVA test) performing a Tukey’s post hoc test to correct for multiple comparisons; when populations were not normally distributed, a Kruskal-Wallis test was used with a Dunn’s post hoc test to correct for multiple comparisons.

Differences among groups are indicated as: *p*-values <0.05 (**\***), *p*-values <0.01 (**\*\***), *p*-values <0.005 (**\*\*\***), *p*-values <0.001 (**\*\*\*\***), and differences among groups not statistically significant (ns).

## Acknowledgements

We thank Prof. Wilfried Weber for generously providing the genetic parts encoding LgBiT and SmBiT, and Dr. Cao Nguyen Duong for assistance with the time-lapse microscopy experiments. We are grateful to Christian Ersfeld for constructing the custom NIR-LED device with culture tube holder used in this study and to Bruno Schäfer for helping to determine its illumination intensity with a power meter. We also acknowledge Varun Sai Tadimarri and Florian Riedel for demonstrating the bead fabrication process, Zirui Ye for assisting with the initial optimization experiments, and Ketaki Deshpande for providing the HUVEC cells. The schematic illustrations were created with BioRender.com. This work was supported by the Leibniz Science Campus (LSC) Living Therapeutic Materials (LifeMat) and DFG Mo2192/4-2.

## 5. Conflict of Interest

The authors declare no conflict of interest.

## 6. Data Availability statement

The information provided in the main text and supporting information are sufficient to reproduce the experiments described in this study. The raw and processed datasets along with relevant metadata are available from the corresponding author upon reasonable request. Received: ((will be filled in by the editorial staff)) Revised: ((will be filled in by the editorial staff)) Published online: ((will be filled in by the editorial staff))

## Table of Contents Entry

**NIR-responsive Engineered Living Materials (ELMs) for controlled angiogenesis:** Near-infrared (800 nm) light activates engineered probiotic bacteria within alginate-based living materials to secrete a blood vessel-regenerating protein. The released protein stimulates endothelial network formation. This platform enables precise, non-invasive, and tunable control of pro-angiogenic signaling for future regenerative medicine applications.

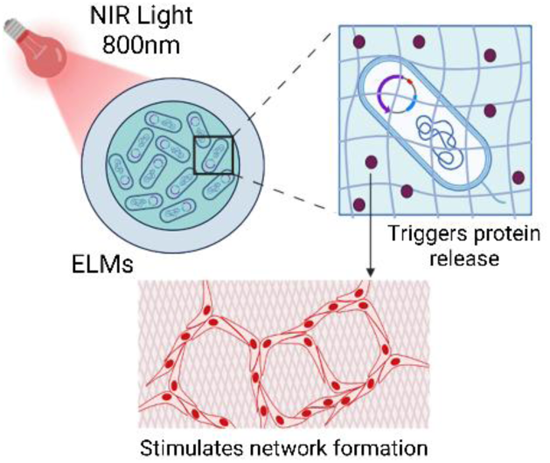

## Notes

### Competing Interest Statement

The authors have declared no competing interest.

## References

1. Stepanov YV, Golovynska I, Golovynskyi S, et al. Red and near infrared light-stimulated angiogenesis mediated via Ca2+ influx, VEGF production and NO synthesis in endothelial cells in macrophage or malignant environments. J Photochem Photobiol B. 2022;227:112388. doi:10.1016/j.jphotobiol.2022.112388

2. Dancáková L, Vasilenko T, Kováč I, et al. Low-level laser therapy with 810 nm wavelength improves skin wound healing in rats with streptozotocin-induced diabetes. Photomed Laser Surg. 2014;32(4):198–204. doi:10.1089/pho.2013.3586

3. Medrado ARAP, Pugliese LS, Reis SRA, Andrade ZA. Influence of low level laser therapy on wound healing and its biological action upon myofibroblasts. Lasers Surg Med. 2003;32(3):239–244. doi:10.1002/lsm.10126

4. Mgwenya TN, Abrahamse H, Houreld NN. Photobiomodulation studies on diabetic wound healing: An insight into the inflammatory pathway in diabetic wound healing. Wound Repair Regen. 2025;33(1):e13239. doi:10.1111/wrr.13239

5. Lee HJ, Kim YK. Burn Wound Successfully Treated with 830-nm Light Emitting Diode Phototherapy Combined with Epidermal Growth Factor Solution. Med Lasers Eng Basic Res Clin Appl. 2019;8(2):94–96. doi:10.25289/ML.2019.8.2.94

6. Carvalho EDS, Rosa RH, de Pereira F M, et al. Effects of diode laser irradiation and fibroblast growth factor on periodontal healing of replanted teeth after extended extra-oral dry time. Dent Traumatol Off Publ Int Assoc Dent Traumatol. 2017;33(2):91–99. doi:10.1111/edt.12308

7. Karpanen T, Bry M, Ollila HM, et al. Overexpression of Vascular Endothelial Growth Factor-B in Mouse Heart Alters Cardiac Lipid Metabolism and Induces Myocardial Hypertrophy. Circ Res. 2008;103(9):1018–1026. doi:10.1161/CIRCRESAHA.108.178459

8. Venkatesan M, Semper C, Skrivergaard S, et al. Recombinant production of growth factors for application in cell culture. iScience. 2022;25(10):105054. doi:10.1016/j.isci.2022.105054

9. Ren X, Zhao M, Lash B, Martino MM, Julier Z. Growth Factor Engineering Strategies for Regenerative Medicine Applications. Front Bioeng Biotechnol. 2020;7. doi:10.3389/fbioe.2019.00469

10. Beheshtizadeh N, Gharibshahian M, Bayati M, et al. Vascular endothelial growth factor (VEGF) delivery approaches in regenerative medicine. Biomed Pharmacother. 2023;166:115301. doi:10.1016/j.biopha.2023.115301

11. Gu Z, Wang J, Fu Y, et al. Smart Biomaterials for Articular Cartilage Repair and Regeneration. Adv Funct Mater. 2023;33(10):2212561. doi:10.1002/adfm.202212561

12. Dong Y, Fu S, Yu J, Li X, Ding B. Emerging Smart Micro/Nanofiber-Based Materials for Next-Generation Wound Dressings. Adv Funct Mater. 2024;34(9):2311199. doi:10.1002/adfm.202311199

13. Nazemidashtarjandi S, Larsen B, Cheng K, et al. Near-infrared light-responsive hydrogels for on-demand dual delivery of proangiogenic growth factors. Acta Biomater. 2024;183:61–73. doi:10.1016/j.actbio.2024.05.052

14. Ma S, Zhang C, Ren X, et al. Photothermally Responsive Hydrogel Releases Basic Fibroblast Growth Factor to Promote the Healing of Infected Wounds. Biomater Res. 29:0156. doi:10.34133/bmr.0156

15. Wang S, Zhang Z, Wei S, et al. Near-infrared light-controllable MXene hydrogel for tunable on-demand release of therapeutic proteins. Acta Biomater. 2021;130:138–148. doi:10.1016/j.actbio.2021.05.027

16. Miao L, Lu X, Wei Y, et al. Near-infrared light-responsive nanocomposite hydrogels loaded with epidermal growth factor for diabetic wound healing. Mater Today Bio. 2025;31:101578. doi:10.1016/j.mtbio.2025.101578

17. Wang L, Wang N, Zhang W, et al. Therapeutic peptides: current applications and future directions. Signal Transduct Target Ther. 2022;7(1):48. doi:10.1038/s41392-022-00904-4

18. Rizzo MG, Palermo N, D’Amora U, et al. Multipotential Role of Growth Factor Mimetic Peptides for Osteochondral Tissue Engineering. Int J Mol Sci. 2022;23(13):7388. doi:10.3390/ijms23137388

19. Ito K, Matsuda Y, Mine A, et al. Single-chain tandem macrocyclic peptides as a scaffold for growth factor and cytokine mimetics. Commun Biol. 2022;5:56. doi:10.1038/s42003-022-03015-6

20. Targeting angiogenesis: Structural characterization and biological properties of a de novo engineered VEGF mimicking peptide | PNAS. Accessed July 16, 2025. https://www.pnas.org/doi/abs/10.1073/pnas.0505047102

21. Diana D, Basile A, Rosa LD, et al. β-Hairpin Peptide That Targets Vascular Endothelial Growth Factor (VEGF) Receptors: DESIGN, NMR CHARACTERIZATION, AND BIOLOGICAL ACTIVITY *. J Biol Chem. 2011;286(48):41680–41691. doi:10.1074/jbc.M111.257402

22. Light-Regulated Pro-Angiogenic Engineered Living Materials - Dhakane - 2023 - Advanced Functional Materials - Wiley Online Library. Accessed July 16, 2025. https://advanced.onlinelibrary.wiley.com/doi/full/10.1002/adfm.202212695

23. Tadimarri VS, Blanch-Asensio M, Deshpande K, et al. PEARL: Protein Eluting Alginate with Recombinant Lactobacilli. Small. n/a(n/a):2408316. doi:10.1002/smll.202408316

24. Meier SS, Hörzing M, Böhm C, et al. Engineering NIR-Sighted Bacteria. eLife. 2025;14. doi:10.7554/eLife.107069.2

25. Downing KJ, McAdam RA, Mizrahi V. *Staphylococcus aureus* nuclease is a useful secretion reporter for mycobacteria. Gene. 1999;239(2):293–299. doi:10.1016/S0378-1119(99)00408-4

26. Fischer AAM, Schatz L, Baaske J, Römer W, Weber W, Thuenauer R. Real-time monitoring of cell surface protein arrival with split luciferases. Traffic. 2023;24(10):453–462. doi:10.1111/tra.12908

27. Han B, Hall FL, Nimni ME. Refolding of a Recombinant Collagen-Targeted TGF-β2 Fusion Protein Expressed in*Escherichia coli*. Protein Expr Purif. 1997;11(2):169–178. doi:10.1006/prep.1997.0784

28. Kleiner-Grote GRM, Risse JM, Friehs K. Secretion of recombinant proteins from E. coli. Eng Life Sci. 2018;18(8):532–550. doi:10.1002/elsc.201700200

29. Gibisch M, Gorecki P, Tauer C, et al. Extracellular peptide production in Escherichia coli by inducible downregulation of lipoprotein Lpp via MicL sRNA. Appl Microbiol Biotechnol. 2025;109(1):136. doi:10.1007/s00253-025-13524-z

30. Emani SS, Kan A, Storms T, et al. Periplasmic stress contributes to a trade-off between protein secretion and cell growth in Escherichia coli Nissle 1917. Synth Biol. 2023;8(1):ysad013. doi:10.1093/synbio/ysad013

31. Kubota Y, Kleinman HK, Martin GR, Lawley TJ. Role of laminin and basement membrane in the morphological differentiation of human endothelial cells into capillary-like structures. J Cell Biol. 1988;107(4):1589–1598. doi:10.1083/jcb.107.4.1589

32. Zudaire E, Gambardella L, Kurcz C, Vermeren S. A Computational Tool for Quantitative Analysis of Vascular Networks. PLOS ONE. 2011;6(11):e27385. doi:10.1371/journal.pone.0027385

33. Nakatsu MN, Sainson RCA, Pérez-del-Pulgar S, et al. VEGF121 and VEGF165 Regulate Blood Vessel Diameter Through Vascular Endothelial Growth Factor Receptor 2 in an in vitro Angiogenesis Model. Lab Invest. 2003;83(12):1873–1885. doi:10.1097/01.LAB.0000107160.81875.33

34. Frykberg RG, Banks J. Challenges in the Treatment of Chronic Wounds. Adv Wound Care. 2015;4(9):560–582. doi:10.1089/wound.2015.0635

35. Colombo E, Signore A, Aicardi S, et al. Experimental and Clinical Applications of Red and Near-Infrared Photobiomodulation on Endothelial Dysfunction: A Review. Biomedicines. 2021;9(3):274. doi:10.3390/biomedicines9030274

36. Meier SSM, Hörzing M, Böhm C, et al. Engineering NIR-Sighted Bacteria. bioRxiv. Preprint posted online August 27, 2025:2025.04.25.650650. doi:10.1101/2025.04.25.650650

37. Stüven B, Stabel R, Ohlendorf R, Beck J, Schubert R, Möglich A. Characterization and engineering of photoactivated adenylyl cyclases. Biol Chem. 2019;400(3):429–441. doi:10.1515/hsz-2018-0375

38. Möglich A. An Open-Source, Cross-Platform Resource for Nonlinear Least-Squares Curve Fitting. J Chem Educ. 2018;95(12):2273–2278. doi:10.1021/acs.jchemed.8b00649

39. Multamäki E, García de Fuentes A, Sieryi O, et al. Optogenetic Control of Bacterial Expression by Red Light. ACS Synth Biol. 2022;11(10):3354–3367. doi:10.1021/acssynbio.2c00259

