## Supplementary information for "An Engineered Living Material with pro-angiogenic activity inducible by near-infrared light"

### An Engineered Living Material to augment Near-Infrared wound therapies with pro-angiogenic activity

**Table S1: Amino acid and DNA sequences of QK-Fusion:**

| Domain Name | Amino Acid Sequence (5'-3') | DNA Sequence (5'-3') |
|---|---|---|
| **ompa signal peptide** | MKKTAIAIAVALAGFATVAQA | atgaaaaaaaccgcgatcgcgatcgccgtggccttagctggtttcgccaccgttgcccaggcg |
| **NucA** | VSTKKLHKEPATLIKAIDGDTVKLMYKGQPMTFRLLLVDTPETKHPKKGVEKYGPEASAFTKKMVENAKKIEVEFDKGQRTDKYGRGLAYIYADGKMVNEALVRQGLAKVAYVYKPNNTHEQHLRKSEAQAKKEKLNIWSEDNADSGQ | agtactaaaaaattacataaagaacctgcgactttaattaaagcgattgatggtgatacggttaaattaatgtacaaaggtcaaccaatgacattcagactattattggttgatacacctgaaacaaagcatcctaaaaaaggtgtagagaaatatggtcctgaagcaagtgcatttacgaaaaaaatggtagaaaatgcaaagaaaattgaagtcgagtttgacaaaggtcaaagaactgataaatatggacgtggcttagcgtatatttatgctgatggaaaaatggtaaacgaagctttagttcgtcaaggcttggctaaagttgcttatgtttacaaacctaacaatacacatgaacaacatttaagaaaaagtgaagcacaagcgaaaaaagagaaattaaatatttggagcgaagacaacgctgattcaggtcaa |
| **SmBit** | VSGWRLFKKIS | gtgagtgggtggcggttgtttaagaagatcagc |
| **CBD** | WREPSFMVLS | tggcgcgaaccgagctttatggtgctgagc |
| **Strep-Tag II** | WSHPQFEK | tggagccacccgcaatttgagaaa |
| **QK** | KLTWQELYQLKYKGID | gatattggcaaatataaactgcagtatctggaacagtggaccctgaaa |

Primers used to create <ins>*Av*</ins>NIRusk- QK-Fusion plasmid:

**NIRusk V fw:** 5'-CACTTTTCGGGGAAATGT-3'

**NIRusk V rev:** 5'-GGTATATCTCCTTCTTAAAGTTAAAC-3'

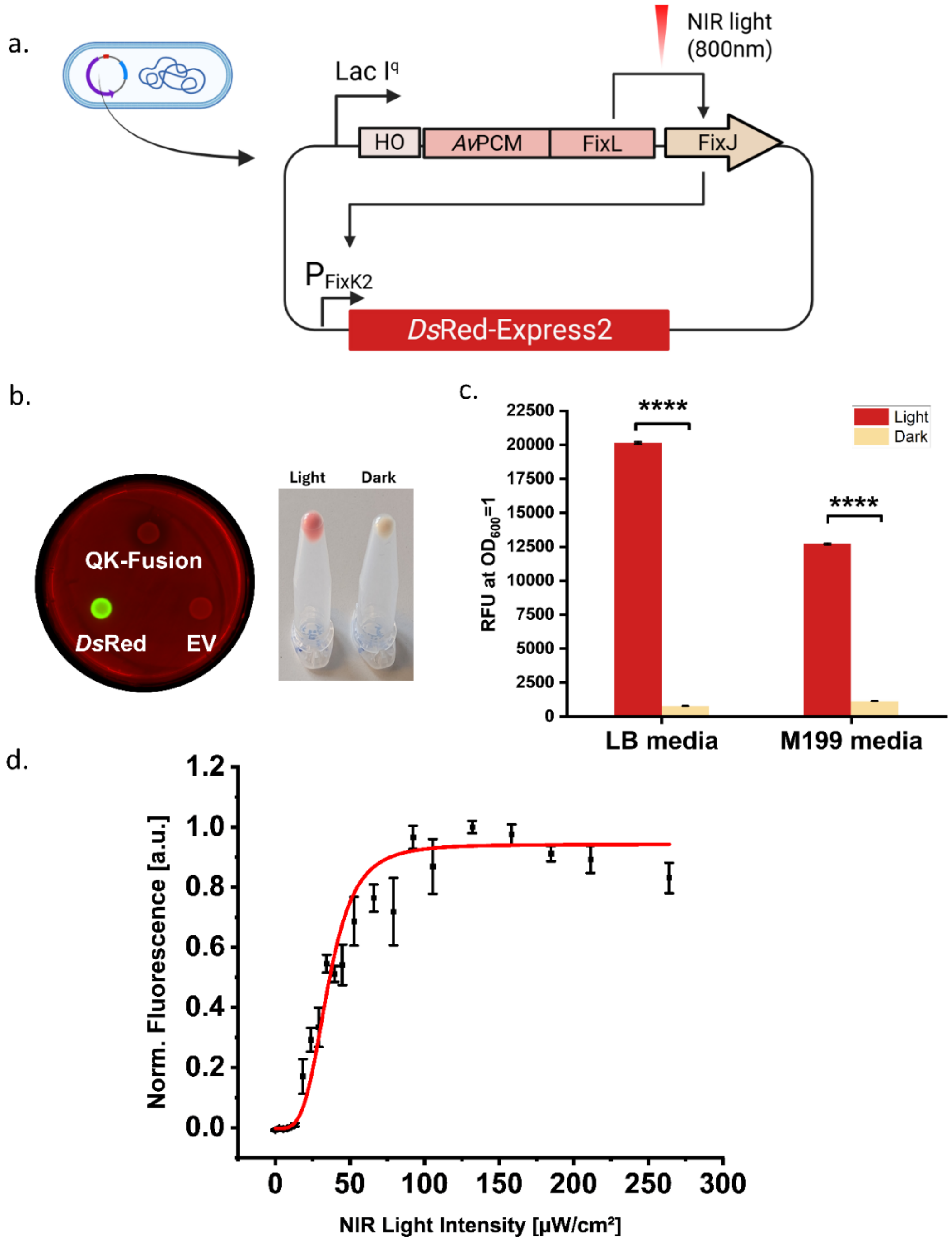


**Fig. S1. Validation of NIR-responsive plasmid system using DsRed reporter**. a. Schematic of the NIR-inducible plasmid construct (pAvNIRusk-DsRed), where DsRed-Express2 is expressed under the control of the PFixK2 promoter. (b) (Left) DNase agar plate supplemented with kanamycin, divided into three sectors and spotted with log phase E. coli Nissle 1917 cultures expressing QK-Fusion, DsRed, or an empty vector (EV). Plates were incubated overnight under NIR light. Fluorescence imaging revealed bright red signal only in the DsRed sector, indicating functional induction. No fluorescence was detected from QK-Fusion or EV. Halo formation, characteristic of

NucA activity, was observed only in the QK-Fusion condition, while DsRed and EV lacked any visible clearing zones. (Right) Visible fluorescence in eppendorf containing cell pellets of overnight cultures of DsRed-expressing EcN under light and dark conditions. Only the light-induced sample showed red coloration, confirming light-responsive expression. (c) Quantification of DsRed fluorescence in LB and M199 media after 16 h induction. Strong induction was observed under light, with negligible background in dark conditions, demonstrating robust and tightly regulated NIR-responsive control across both media types. Data shown as mean ± s.d.; ****p < 0.0001 (N,n = 3). (d) Dose–response curve showing normalized DsRed fluorescence in *E. coli* Nissle 1917 expressing the AvNIRusk circuit when cultured in LB medium and exposed to increasing 800 nm NIR light intensities (0–250 μW $cm^{-2}$). Fluorescence increases steeply from 20 μW $cm^{-2}$ and reaches a plateau at 80 μW $cm^{-2}$, indicating saturation of circuit activation. Data are presented as mean ± s.d. (N,n=3). Fitting curve: Hill isotherm

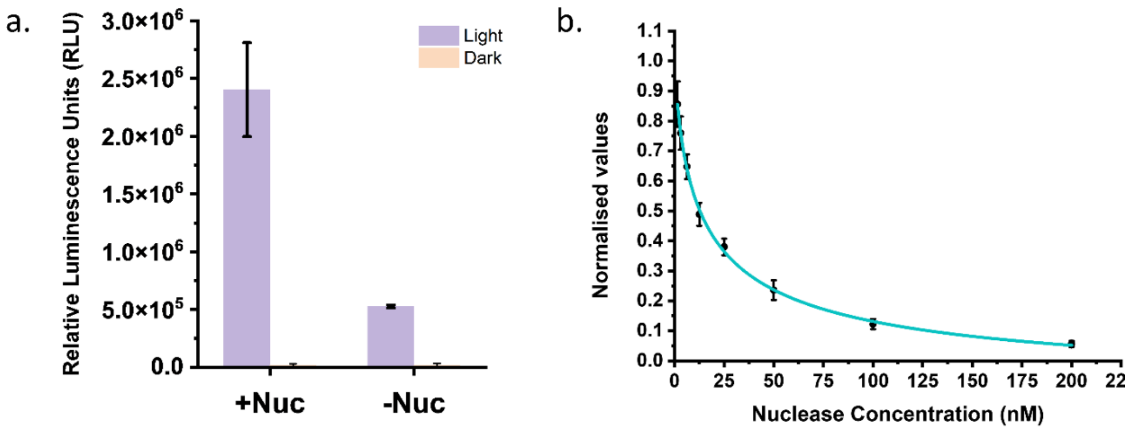


**Fig. S2. Role of the NucA domain in secretion quantification.** (a) Split-luciferase assay comparing supernatants from NIR-induced (+Light) and non-induced (Dark) conditions for strains expressing QK-Fusion either with the NucA domain (+Nuc) or without it (−Nuc). The presence of NucA enables strong luminescent signal under NIR induction, whereas constructs lacking NucA produce significantly lower signal, confirming that NucA improves secretion efficiency manifolds. Data shown as mean ± s.d. (N,n = 3). (b) Standard curve from known concentrations of purified NucA protein (1.56 nM, 3.12 nM, 6.25 nM, 12.5 nM, 25 nM, 50 nM, 100nM, 200nM). Normalised values were fitted to an exponential decay function and used to convert experimental fluorescence measurements to protein concentration. (N=2, n=3)

a.

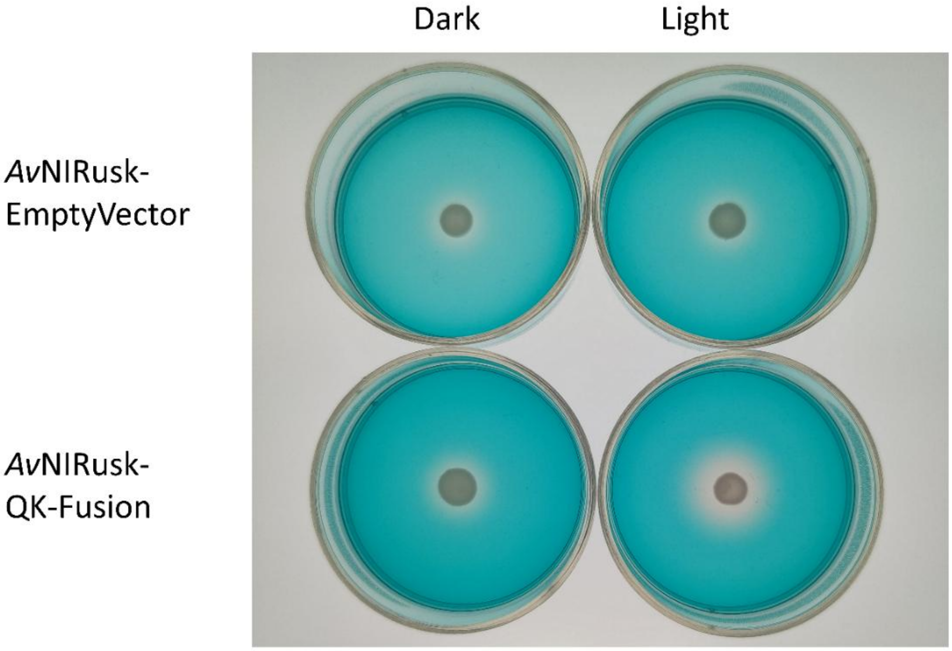


b.

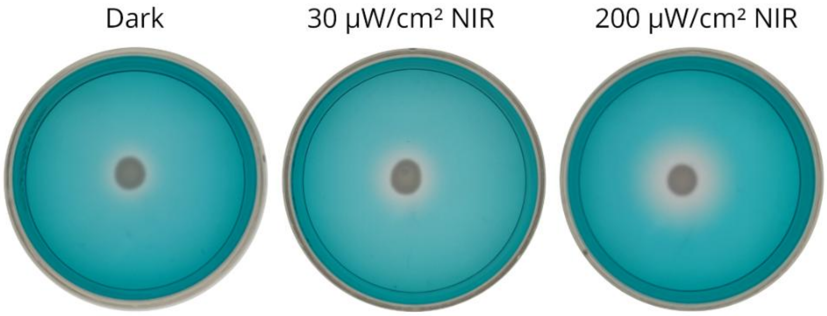


**Fig. S3:** Nuclease Assay on DNAse-agar plates confirming light-dependent secretion and intensity-dependent tuning. (a) DNase agar plates spotted with supernatants from *E. coli* Nissle 1917 expressing either the AvNIRusk–Empty Vector or the AvNIRusk–QK-Fusion construct, incubated under Dark or NIR Light conditions. Clear halos indicating DNase activity were observed only for the QK-Fusion strain under NIR induction, confirming light-dependent secretion of the fusion protein. (b) DNase agar plates demonstrating dose-dependent secretion, where increasing NIR intensities (0, 30, and 200 µW $cm^{-2}$) resulted in progressively larger halo diameters for the QK-Fusion strain. This confirms that secretion levels can be tuned by varying NIR illumination intensity.

| Trial ID | Core alginate Concentration (% w/v) | $CaCl_2$ Concentration (wt%) for core | Core Crosslinking Time (min) | Shell alginate Concentration (% w/v) | $CaCl_2$ Concentration (wt%) for shell | Shell Crosslinking Time (hours) | Medium Used in Core | Medium Used in Surrounding | Bacterial Growth Phase Before Encapsulation | Bead incubation temperature (°C) | Containment (Day of Leakage) | Growth within Core | Protein Release (Qualitative) |
|---|---|---|---|---|---|---|---|---|---|---|---|---|---|
| 1 | 2 | 5 | 30 | 1,5 | 5 | 0,5 | LB | LB | Mid-log | 37 | 1 | Good | N/A |
| 2 | 2 | 5 | 30 | 1,5 | 5 | 1 | LB | LB | Mid-log | 37 | 1 | Good | N/A |
| 3 | 2 | 5 | 30 | 1,5 | 5 | 2 | LB | LB | Mid-log | 37 | 1 | Good | N/A |
| 4 | 2 | 5 | 30 | 1,5 | 5 | 3 | LB | LB | Mid-log | 37 | 1 | Good | N/A |
| 5 | 2 | 5 | 30 | 1,5 | 5 | 4 | LB | LB | Mid-log | 37 | 1 | Good | N/A |
| 6 | 2 | 6 | 30 | 1,5 | 6 | 4 | LB | LB | Mid-log | 37 | 1 | Good | N/A |
| 7 | 2 | 5 | 30 | 1,5 | 5 | 4 | LB | M199 | Mid-log | 37 | 1 | Mid | N/A |
| 8 | 2 | 5 | 30 | 2 | 5 | 4 | LB | M199 | Mid-log | 37 | 2 | Mid | Negligible |
| 9 | 2 | 5 | 30 | 1,5 | 6 | 4 | LB | M199 | Mid-log | 37 | 2 | Mid | Low |
| 10 | 2 | 6 | 30 | 1,5 | 6 | 4 | LB | M199 | Mid-log | 37 | 2 | Mid | Low |
| 11 | 2 | 6 | 30 | 2 | 6 | 4 | LB | M199 | Mid-log | 37 | 2 | Mid | Low |
| 12 | 2 | 5 | 30 | 1,5 | 5 | 0,5 | M199 | M199 | Mid-log | 37 | 2 | Sparse | Low |
| 13 | 2 | 5 | 30 | 1,5 | 5 | 1 | M199 | M199 | Mid-log | 37 | 2 | Sparse | Low |
| 14 | 2 | 5 | 30 | 1,5 | 5 | 2 | M199 | M199 | Mid-log | 37 | 4 | Sparse | Low |
| 15 | 2 | 5 | 30 | 1,5 | 5 | 3 | M199 | M199 | Mid-log | 37 | 10 | Sparse | Low |
| 16 | 2 | 6 | 30 | 1,5 | 6 | 4 | M199 | M199 | Mid-log | 37 | 10 | Sparse | Low |
| 17 | 2 | 6 | 30 | 1,5 | 6 | 4 | M199 + 0,4% w/v Glucose | M199 | Mid-log | 37 | 3 | Good | Low |
| 18 | 2 | 6 | 30 | 2 | 6 | 4 | M199 + 0,4% w/v Glucose | M199 | Mid-log | 37 | 4 | Good | Low |
| 19 | 2 | 6 | 30 | 2,5 | 6 | 4 | M199 + 0,4% w/v Glucose | M199 | Mid-log | 37 | 7 | Good | Negligible |
| 20 | 2 | 8 | 30 | 1,5 | 8 | 4 | M199 + 0,4% w/v Glucose | M199 | Mid-log | 37 | 10 | Good | Low |
| 21 | 2 | 8 | 30 | 1,5 | 10 | 4 | M199 + 0,4% w/v Glucose | M199 | Mid-log | 37 | 10 | Good | Negligible |
| 22 | 2 | 10 | 30 | 1,5 | 10 | 4 | M199 + 0,4% w/v Glucose | M199 | Mid-log | 37 | 10 | Good | Negligible |
| 23 | 2 | 6 | 30 | 1,5 | 8 | 4 | M199 + 0,4% w/v Glucose | M199 | Mid-log | 37 | 3 | Good | High |
| 24 | 3 | 6 | 30 | 2 | 8 | 4 | M199 + 0,4% w/v Glucose | M199 | Mid-log | 37 | 7 | Mid | Low |
| 25 | 3 | 6 | 30 | 1,5 | 8 | 4 | M199 + 0,4% w/v Glucose | M200 | Mid-log | 38 | 7 | Good | Negligible |
| 26 | 2 | 6 | 30 | 3 | 8 | 4 | M199 + 0,4% w/v Glucose | M199 | Mid-log | 37 | 7 | Good | Negligible |
| 27 | 2 | 6 | 30 | 1,5 | 8 | 4 | M199 + 0,4% w/v Glucose | M199 | late log | 37 | 3 | Good | Low |
| 28 | 2 | 6 | 30 | 1,5 | 8 | 4 | M199 + 0,4% w/v Glucose | M199 | Early log | 37 | 3 | Good | Very High |
| 29 | 2 | 6 | 30 | 1,5 | 8 | 4 | M199 + 0,4% w/v Glucose | M199 | Early log | 30 | 10 | Good | Very High |

**Table S2:** Iterative optimization of encapsulation parameters for developing a stable core–shell Engineered Living Material. Summary of encapsulation trials performed to identify the optimal conditions for bacterial containment and protein secretion from *E. coli* Nissle 1917 encapsulated in alginate beads.

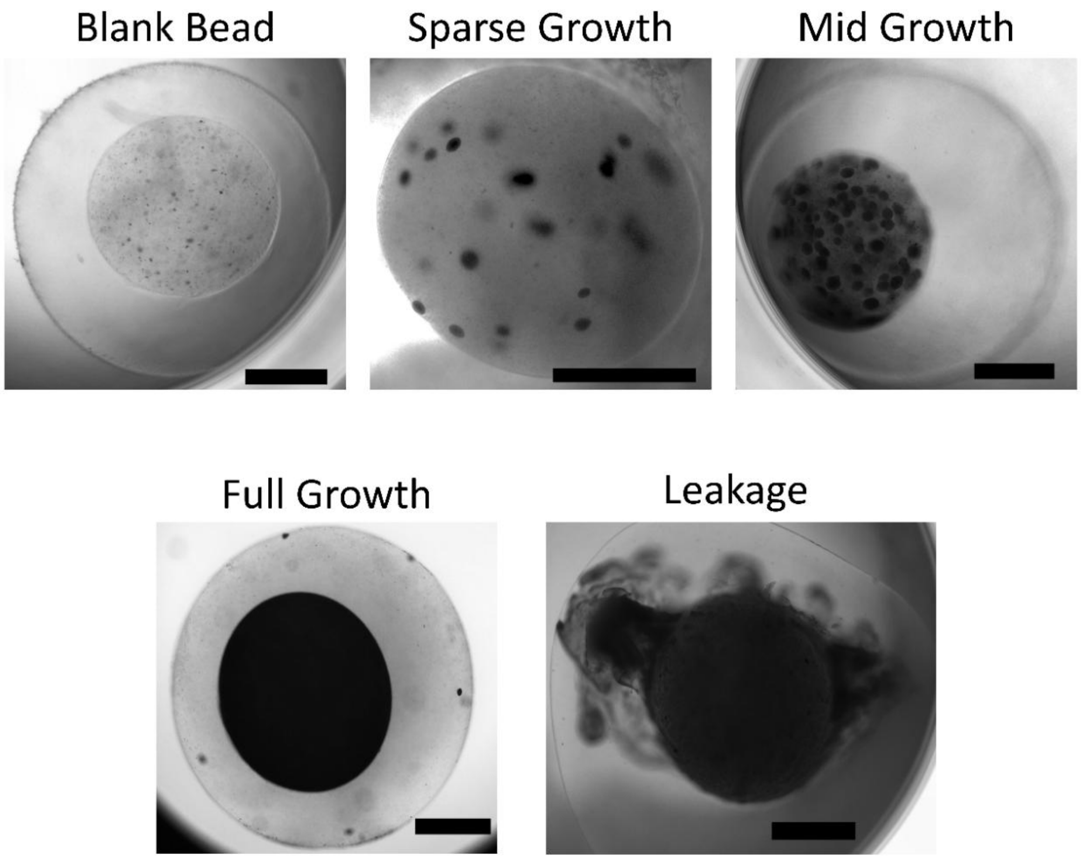


**Fig. S4:** Representative brightfield images showing different bacterial growth behaviour in alginate core-shell beads. Images illustrate (top row, left to right) a blank bead with no bacteria, a bead showing *sparse growth* with few visible microcolonies, and a bead with *mid growth* where more bacterial colonies have grown. (Bottom row) *Full growth* indicates uniform bacterial proliferation throughout the core, while *leakage* represents uncontrolled bacterial escape and overgrowth outside the bead boundary. All images were taken after 24 hours of encapsulation. Scale bars: 1 mm.

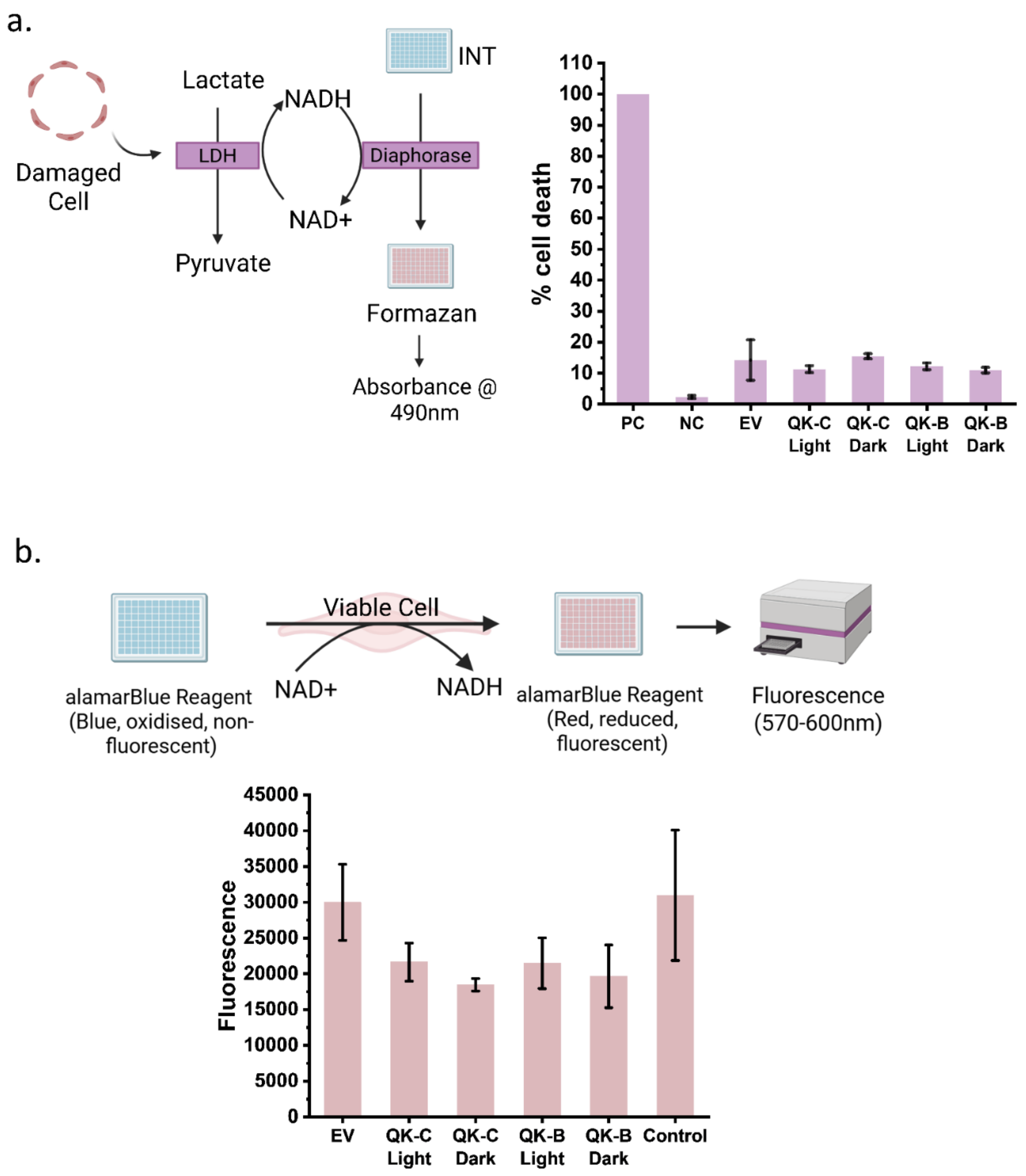


**Fig. S5:** Cytocompatibility assessment of ELMs and bacterial cultures on HUVECs. (a) (Left) Schematic of Lactate Dehydrogenase Assay. (Right) LDH assay showing percent cell death in HUVECs treated with supernatants from QK-Fusion producing bacterial cultures (QK-C) and beads (QK-B) under light and dark conditions, compared to positive control (PC), negative control (NC), and empty vector (EV). (b) (Top) Schematic of AlamarBlue Assay. (Bottom) AlamarBlue assay showing metabolic activity of HUVECs treated with supernatants from QK-Fusion producing bacterial cultures (QK-C) and beads (QK-B) under light and dark conditions, compared to growth medium as control and empty vector (EV). Both assays indicate no significant cytotoxicity from supernatants. Graphs are shown as data ± S.D. (N=3, n ≥ 3)

#### **Supplementary Code S1. Fiji macro for batch image analysis of angiogenic networks**

This macro automates batch image processing and segmentation of brightfield microscopy images from HUVEC Matrigel assays. It was used to generate the quantitative data shown in Figures 5c–f. The script sequentially processes all image files in a selected directory, applies a pre-trained *Trainable Weka Segmentation* classifier, thresholds the probability maps, performs morphological clean-up, and exports binary masks suitable for *AngioTool* analysis.

Input: Raw images (TIFF format).
Output: Binary masks (TIFF).
Software: *Fiji* (ImageJ, NIH) with *Trainable Weka Segmentation* v3.3.3.

```
#@ File(label="Input directory", description="Select the directory with input .lof images",
style="directory") inputDir
#@ File(label="Output directory", description="Select the output directory for binary masks",
style="directory") outputDir
#@ File(label="Weka model", description="Select the Weka model to apply") modelPath
#@ String(label="Result mode",choices={"Labels","Probabilities"}, style="radioButtonHorizontal")
resultMode

import trainableSegmentation.WekaSegmentation
import trainableSegmentation.utils.Utils
import ij.io.FileSaver
import ij.IJ
import ij.ImagePlus

long startTime = System.currentTimeMillis()
boolean getProbs = resultMode.equals("Probabilities")

// Create segmentator and load classifier once
WekaSegmentation segmentator = new WekaSegmentation()
segmentator.loadClassifier(modelPath.getCanonicalPath())

// List files in input directory
File[] listOfFiles = inputDir.listFiles()

for (File file : listOfFiles)
 {
   if (file.isFile() && file.getName().toLowerCase().endsWith(".lof"))
         {
               IJ.log("Processing: " + file.getName())

               ImagePlus image = IJ.openImage(file.getCanonicalPath())
               if (image == null)
                    {
                          IJ.log("Could not open image: " + file.getName())
                          continue
                    }
```

```
        // Apply classifier on image
        ImagePlus result = segmentator.applyClassifier(image, 0, getProbs)
        if (!getProbs)
            {
                // Apply LUT like GUI does (optional)
                result.setLut(Utils.getGoldenAngleLUT())
            }

        // Post-processing: threshold + mask + max filter + skeletonize
        IJ.setThreshold(result, 1, 2)
        IJ.run(result, "Convert to Mask", "")
        IJ.setAutoThreshold(result, "Default dark")
        IJ.run(result, "Convert to Mask", "")


        // Save binary mask as TIFF
            def baseName = file.getName().replaceFirst(/[.][^.]+$/, "")
                          def outPath = outputDir.getPath() + File.separator + baseName + ".tif"
                          new FileSaver(result).saveAsTiff(outPath)


                // Clean up
                image.close()
                result.close()
                image = null
                result = null
                System.gc()
            }
}

long elapsedTime = System.currentTimeMillis() - startTime
IJ.log("** Finished processing folder in " + elapsedTime + " ms **")
```
